# Anti-apoptotic BH3-only proteins inhibit Bak-dependent apoptosis

**DOI:** 10.1101/2022.07.24.499430

**Authors:** Sebastian Ruehl, Clifford S. Guy, Zhenrui Li, Mao Yang, Tudor Moldoveanu, Douglas R. Green

## Abstract

Bcl-2 family proteins regulate induction of intrinsic apoptosis through initiating mitochondrial outer membrane permeabilization (MOMP). Activation of the MOMP effectors Bax and Bak is controlled by interplay levels of anti-apoptotic Bcl-2 proteins (e.g. Mcl-1) and pro-apoptotic BH3-only proteins (e.g. BIM). Using a genome-wide CRISPR-dCas9 transactivation screen we identified two Bcl-2 family proteins, BNIP5 and Bcl-G, as inhibitors of Bak, but not Bax induced apoptosis. BNIP5 was able to block Bak activation in different cell types and in response to various cytotoxic therapies. The BH3 domain of BNIP5 was both necessary and sufficient to block Bak activation. Mechanistically, the BH3 domains of BNIP5 and Bcl-G act as a selective Bak activators, while not inhibiting anti-apoptotic proteins. This led to increased binding of activated Bak to Mcl-1, which prevented apoptosis engagement, identifying BNIP5 and Bcl-G as anti-apoptotic BH3-only proteins.

## Introduction

The Bcl-2 family of proteins (Bcl-2 proteins) are at the core of the mitochondrial pathway of apoptosis(Chipuk et al., 2010). This protein family regulates the crucial event of mitochondrial outer membrane permeabilization (MOMP), which commits cells to the execution of apoptosis(Kale et al., 2018). Bcl-2 proteins harbor varying numbers of Bcl-2 homology domains (BH domains), numbered BH1-BH4. These BH domains are short linear motifs of 18-24 amino acids, and the BH3 domain, present in all Bcl-2 proteins, is a crucial determinant of the interaction specificity between the different members of the family. The ‘modes’ of interaction between these proteins are summarized in a ‘unified model’ (Llambi et al., 2011). The two MOMP effectors Bcl-2 associated x (Bax) and Bcl-2 antagonist killer (Bak) are inactive in resting cells as these cells exhibit low levels of proapoptotic BH3-only proteins (e.g. BIM). These low levels of BH3-only proteins can be bound and neutralized by multidomain anti-apoptotic proteins (e.g. Bcl-xL). This state is referred to as ‘MODE 1’ inhibition. All known BH3-only proteins exert their proapoptotic function through inhibition of anti-apoptotic Bcl-2 proteins (sensitizers, e.g. BAD) and some are additionally able to activate Bax and Bak (sensitizers and direct activators, e.g. BIM).

When apoptotic stress increases, for example through treatment with chemotherapeutic drugs, the effectors Bax and Bak are activated by rising levels of BH3-only proteins and the activated effectors are sequestered by anti-apoptotic proteins to form ‘MODE 2’ complexes(Llambi et al., 2011). Further increase of apoptotic stress overwhelms the ability of anti-apoptotic proteins to inhibit Bax and Bak, which are then able to oligomerize and permeate the outer mitochondrial membrane (OMM) to promote cytochrome c release and apoptosis (Chen et al., 2015; Llambi et al., 2011). While the BH3 domains of BH3-only proteins are sufficient to execute their sensitization or activation function(Letai et al., 2002), interaction of activated Bak with anti-apoptotic proteins requires proper assembly of the BH groove of the anti-apoptotic proteins, including BH1 and BH2 domains (Zheng et al., 2016). The stepwise activation of Bak from dormant to active has been structurally characterized (Birkinshaw et al., 2021; Brouwer et al., 2014; Moldoveanu et al., 2013; Sandow et al., 2021) and we know that Bak exposes its BH3 domain upon activation (Dai et al., 2011), which can either bind into the BH groove of anti-apoptotic proteins to form MODE 2 complexes or bind to activate other Bax and Bak molecules(Czabotar et al., 2013; Iyer et al., 2020; Singh et al., 2022) to induce pore formation through oligomerization (Cowan et al., 2020). Structure-guided drug design efforts to mimic the function of BH3-only sensitizer proteins led to the development of BH3 mimetics, which act as selective inhibitors of the anti-apoptotic Bcl-2 proteins (Montero and Letai, 2018).

Besides the functions described above, anti-apoptotic Bcl-2 proteins can also function to shuttle Bax and Bak between the MOM and the cytosol by a process called ‘retro-translocation’(Edlich et al., 2011; Xu et al., 2011). Translocation rates for Bax are faster than for Bak, as the latter is stably integrated into the MOM through its C-terminal transmembrane domain(Todt et al., 2015). Removal of either effector from the MOM by anti-apoptotic proteins contributes to the inhibitory function of these proteins.

Both Bax and Bak can induce MOMP, and despite minor differences in their binding specificities for anti-apoptotic proteins and BH3-only proteins, are thought to be functionally redundant(Sarosiek et al., 2013). Only deletion of both, Bax and Bak renders cells resistant to most chemotherapeutics(Lindsten et al., 2000) and severely disrupts embryonic development(Ke et al., 2018).

In this study we used an unbiased CRISPR-Cas9 transactivation screen to find novel regulators of Bax- or Bak-induced apoptosis to better understand the mechanisms governing the regulation of each effector. We found that two Bcl-2 proteins, Bcl-2 interacting protein 5 (BNIP5) and Bcl-G, act as selective inhibitors of Bak-dependent but not Bax-dependent apoptosis. The BH3 domains of BNIP5 and Bcl-G were both necessary and sufficient to inhibit Bak-dependent apoptosis. Expression of BNIP5 resulted in stable removal of Bak from mitochondria through the E3 ligase MARCH5 and a proteasome-dependent mechanism. This removal was dispensable for BNIP5-mediated inhibition of Bak. Mechanistically, the BH3 domains of BNIP5 and Bcl-G act as selective Bak activators which increase formation of Mcl-1-Bak complexes. Formation of these complexes, but not Bak-Bcl-xL complexes was required for inhibition of Bak dependent apoptosis by BNIP5.

## Results

### A CRISPR-dCas9 transactivation screen uncovers novel Bcl-2 proteins as Bak inhibitors

To uncover human genes whose products inhibit BH3 mimetic-induced apoptosis we used an unbiased genome-wide CRISPR-dCas9 transactivation (CRISPRa) screen. In CRISPRa screens single guide RNAs (sgRNAs) target a catalytically inactive Cas9-VP64 (dCas9) to the promoter region of a gene of interest. The dCas9-VP64 fusion along with artificial MS2 stem loops in the tracrRNA recruits potent transcriptional activators to express the targeted gene(Sanson et al., 2018). Wildtype, Bax-deficient or Bak-deficient dCas9-VP64-expressing HeLa cells were transduced with the human Broad GPP activation Calabrese p65-HSF library, consisting of approximately 110,000 sgRNA targeting more than 18,000 genes. Cells were selected for 7 days in puromycin and subjected to three rounds of selection with the BH3 mimetics ABT-737 and S63845, which combined inhibit the pro-survival BCL-2 family proteins BCL-2, BCL-xL, BCL-w and MCL-1 (Figure 1A). Enrichment analysis of sgRNAs after Next-Generation Sequencing revealed that in wildtype and Bak-deficient HeLa cells, only upregulation of the pro-survival BCL-2 family protein Bcl2A1 by CRISPRa conferred protection against BH3 mimetic treatment (Figure 1B and C).

**Figure 1:**
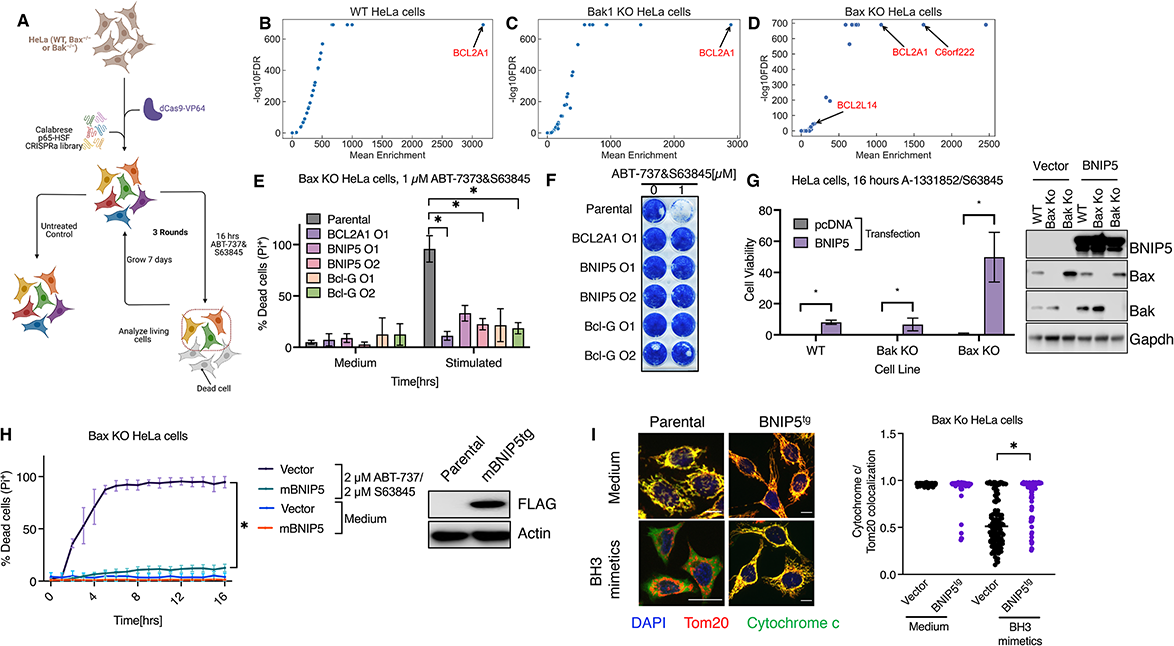
A CRISPR-dCas9 transactivation screen uncovers novel Bcl-2 proteins as Bak inhibitors. **A** Schematic representation of genome-wide CRISPR transactivation (CRISPRa) screen in HeLa cells **B-D** Gene enrichment vs. −log_10_ false-discovery rate (FDR) of CRISPRa screen in wildtype(**B**), Bak1 KO (**C**) or Bax KO HeLa cells (**D**) treated as described in **A**. Genes, where more than 3 sgRNAS were recovered are shown in red. **E-F** HeLa dCas9-VP64 expressing, Bax KO cells expressing indicated sgRNAs were stimulated with 1 µM ABT-737 and 1 µM S63845 and PI positive cells (**E**) or colony formation (**F**) was analyzed after 16 hours or 7 days respectively. **G** Wildtype, Bax or Bak deficient HeLa cells were transfected with either pcDNA or HA-BNIP5 and subsequently stimulated with 1 µM A-1331852 and 1 µM S63845. Cell viability was determined 16 hours post-stimulation. Western Blot panel shows expression of indicated proteins at 24 hours after transfection. **H** Bax deficient HeLa cells expressing murine FLAG-BNIP5 or an empty vector were stimulated with 2 µM ABT-737 and S63845 and PI incorporation was measured over 16 hours. Expression of BNIP5 was analyzed by western blot. **I** Bax deficient HeLa cells, expressing BNIP5 by CRISPRa were stimulated for 2 hours with 1 µM A-1331852 and S63845 in the presence of 40 µM Q-VD-OPh and stained for Tom20, cytochrome C or DNA (DAPI). Colocalization of cytochrome with Tom20 was quantified for all conditions. * p<0.05. All panels except **F** show data of at least 3 pooled independent biological experiments. **F** and western blots are representative of at least 2 independent biological experiments.

In contrast, in Bax-deficient HeLa cells, in addition to the upregulation of Bcl2A1, sgRNAs targeting the promoter region of *C6orf222* and *BCL2L14* were also strongly overrepresented in cells recovered after BH3 mimetic stimulation (Figure 1D). *C6orf222* encodes a 72 kDa Bcl-2 family protein called Bcl-2 interacting protein 5 (BNIP5) and *BCL2L14* encodes the Bcl-2 family member Bcl-G. We validated these results by expressing sgRNAs targeting the promoter region of Bcl2A1, BNIP5 or Bcl-G in dCas9-VP64-expressing Bax-deficient HeLa cells and assessed cell death after treatment with BH3 mimetics using propidium iodide (PI) incorporation. Cells overexpressing BNIP5, like cells expressing Bcl2A1 or Bcl-G, were significantly protected against BH3 mimetic-induced loss of cell viability as assessed by PI incorporation and displayed increased clonogenic survival (Figure 1E and 1F). Western blot analysis revealed the mutually exclusive expression of Bcl2A1, BNIP5, or Bcl-G in the respective CRIPSRa cell lines, suggesting that the proteins block Bak-induced apoptosis independently of each other and that they are not expressed at detectable level in the parental line (Figure S1A).

We further asked if upregulation of BNIP5 or Bcl-G by CRISPRa in other cell lines also protected from Bak induced apoptosis. We transduced PC9 lung cancer or A375 melanoma cells with dCas9-VP64 and either a control sgRNA or sgRNAs targeting the promoter regions of BNIP5 or Bcl-G. Subsequently, cells were transfected with a control siRNA or siRNAs targeting either Bax or Bak. Only if Bax expression was reduced by siRNA transfection, and hence cells were primarily dependent on Bak for apoptosis in response to BH3 mimetic treatment, BNIP5 or Bcl-G expression protected these cells from apoptosis (Figure S1B-1G). We note the expression of BNIP5 or Bcl-G decreased the level of Bak compared to untreated control and we will investigate this observation below (Figure S1C, S1D, S1F, S1G).

Because of technical difficulties in expressing the Bcl-G cDNA, we focused our efforts on the investigation of BNIP5. Transient expression of BNIP5 in wildtype, Bak-or Bax-deficient HeLa cells confirmed that Bax-deficient HeLa cells were most strongly protected from BH3 mimetic-induced apoptosis upon BNIP5 expression (Figure 1G). Heterologous complementation experiments revealed that murine and human BNIP5, expressed in Bax-deficient immortalized murine embryonic fibroblasts (MEFs), significantly reduced BH3 mimetic-induced apoptosis (Figure S1H). Expression of murine BNIP5 in Bax-deficient HeLa cells completely protected these cells from BH3 mimetic-induced apoptosis (Figure 1H). This suggests an evolutionary conserved mechanism of BNIP5 to inhibit Bak-induced apoptosis.

Upon activation, Bak oligomerizes to permeabilize the OMM and release cytochrome c into the cytosol. We assessed MOMP in the presence and absence of BNIP5 in Bax-deficient HeLa cells. Cytochrome c did not localize to mitochondria upon treatment of Bax-deficient HeLa cells with BH3 mimetics but was retained in mitochondria in BNIP5-expressing cells (Figure 1I). Therefore, BNIP5 or Bcl-G expression inhibits Bak-induced apoptosis and promotes long term survival upon BH3 mimetic treatment in a variety of cell types.

### The BH3 domain of BNIP5 is necessary and sufficient to inhibit Bak activation in cells

BNIP5 contains two BH3 domain-like motifs, one in the center, and one at the C-terminus of the protein (Figure 2A). BH3 domains are crucial for the interactions between Bcl-2 proteins and partially determine the specificity of Bcl-2 proteins toward each other. This is particularly true for BH3-only proteins (Kale et al., 2018). BH3 domains contain 3-4 hydrophobic residues which allow binding of the BH3 domain into the BH hydrophobic pockets of anti-apoptotic and effector Bcl-2 proteins, where the BH3 engages in hydrophobic interactions with the corresponding residues of the interaction partner(Aouacheria et al., 2015; Shamas-Din et al., 2013, 2011). To test if any of the two BH3 domains is required to inhibit Bak-dependent apoptosis, we truncated BNIP5 from the N-terminus or C-terminus, transiently expressed these mutants in Bax-deficient HeLa cells, and measured cell viability upon treatment with BH3 mimetics. No drastic loss of cell viability was observed upon expression of any of the truncated mutants (Figure 2B). All of these mutants contain the BH3 domain in the center of BNIP5 but mutants C2 and C3 lack the C-terminal BH3 motif (Figure 2A). This suggests that the C-terminal BH3 domain is dispensable for Bak inhibition by BNIP5. A genome-wide computational prediction of BH3 domains identified the central BH3 domain (D253-S278) as a canonical BH3 domain (DeBartolo et al., 2014). Deletion of these 24 amino acids (ΔBH3) or mutation of the four hydrophobic residues to arginine (4R) completely abolished the ability of BNIP5 to inhibit cell death (Figure 2B), while all mutants tested were expressed at comparable levels (Figure S2A).

**Figure 2:**
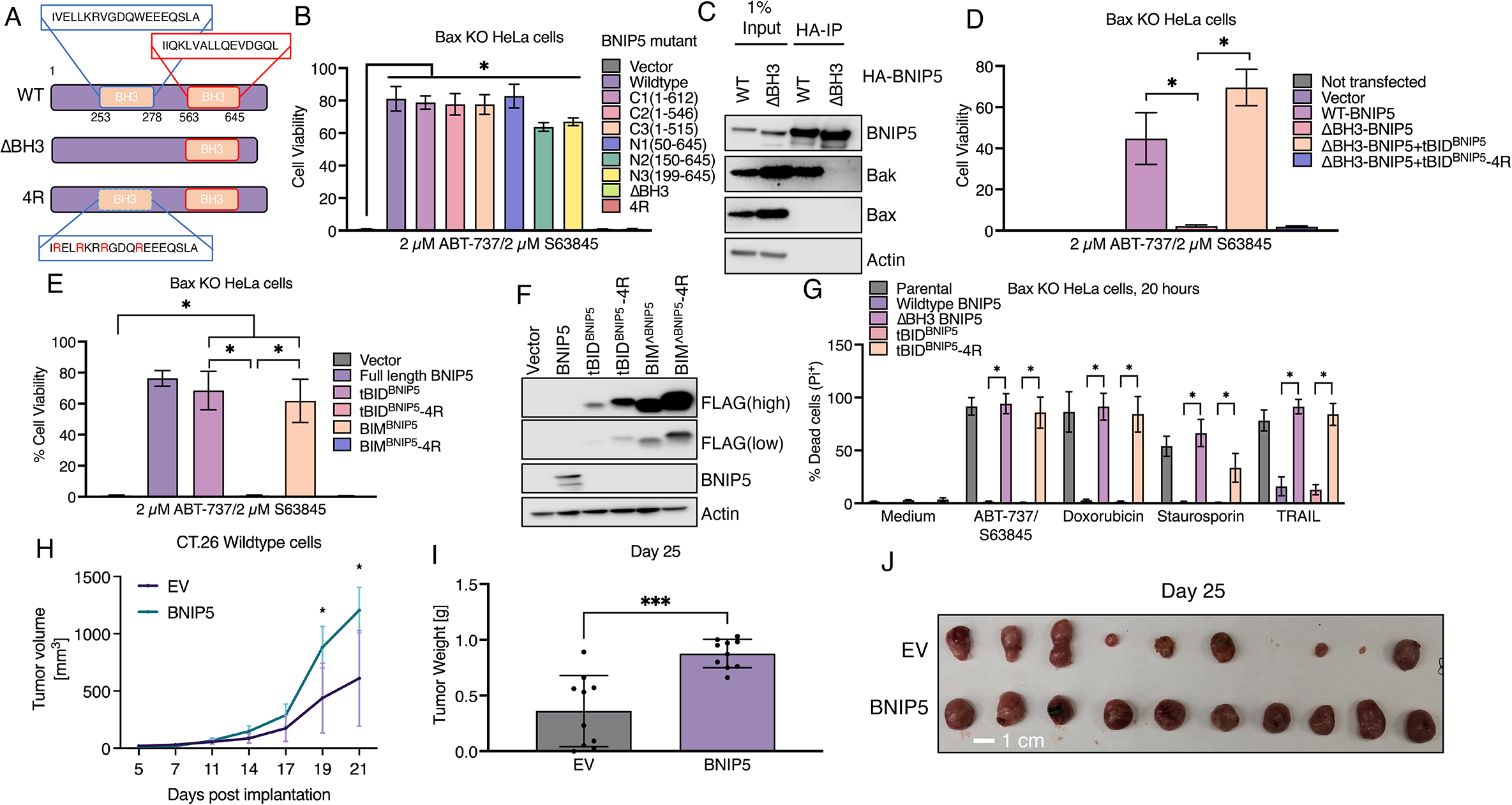
The BH3 domain of BNIP5 is necessary and sufficient to inhibit Bak activation in cells. **A** Schematic representation of BNIP5 mutants used in Figure 2B and **S2B**. **B**. ATP-based cell viability assay of Bax deficient HeLa cells transfected with indicated BNIP5 mutants and subsequently stimulated with 2 µM ABT-737&S63845. **C** HA-BNIP5 immunoprecipitation from transiently transfected HEK 293T cells, 48 hours post transfection. **D-E** ATP-based cell viability assay of Bax deficient HeLa cells transfected with the indicated combinations of plasmids and treated with 2 µM ABT-737&S63845. **F** Western blot shows expression of transfected constructs in **E**. **G** Bax deficient HeLa cells were stably transduced with indicated constructs and stimulated with 2 µM ABT-737&S63845, 1 µM Doxorubicin, 250 nM staurosporin or 100 ng/ml recombinant TRAIL for 24 hours. Number of PI positive cells were enumerated using an Incucyte imaging system. **H-J** Tumor growth curves (**H**), weights (**I**), and pictures of tumors (**J**) of wildtype CT.26 cells expressing either an empty vector (EV) or BNIP5 implanted into Balb/c mice. * p<0.05 **A**,**C**,**D**,**F** show pooled data from 3 independent biological experiments. Western blots are representative of at least 2 independent biological replicates.

We hypothesized that the BH3 domain of BNIP5 mediates interaction with Bak to allow inhibition of Bak-dependent apoptosis. Immunoprecipitation experiments in transiently transfected HEK293T cells showed that endogenous Bak co-precipitated with HA-tagged wildtype BNIP5, but not ΔBH3 BNIP5, while Bax did not precipitate with either BNIP5 construct (Figure 2C). These results suggest that BNIP5 interacts with Bak via its BH3 domain to inhibit apoptosis.

To interrogate if expression of the BNIP5 BH3 domain *in trans* is sufficient to block Bak induced apoptosis, we transfected cells with either WT BNIP5 or ΔBH3, together with a tBID chimera(Hockings et al., 2018, 2015; Llambi et al., 2011) containing the BNIP5 BH3 domain (tBID^BNIP5^) or an inactive mutant (tBID^BNIP5^-4R) in place of the tBID BH3. This allows for expression and correct folding of different BH3 domains in cells. Co-expression of tBID^BNIP5^, but not tBID^BNIP5^-4R with ΔBH3 BNIP5 restored cell viability as measured by ATP based cell viability in response to BH3 mimetic treatment (Figure 2D). To test whether this BH3 domain is sufficient to block Bak-dependent apoptosis, we replaced the BH3 domain of murine BIM-short (BIMs) with the BNIP5 BH3 domain and transfected Bax-deficient HeLa cells with the tBID^BNIP5^, BIMs^BNIP5^ or the inactive BH3 mutant chimeras. Cells expressing tBID^BNIP5^ or BIMs^BNIP5^ were protected from BH3 mimetic-induced apoptosis, while the respective BH3 domain mutants (4R) did not protect (Figure 2E), although the inactive mutants were expressed at higher levels (Figure 2F).

Although we were unable to express full length Bcl-G to recapitulate the results obtained in CRISPRa cells, we noted significant homology between the BNIP5 BH3 domain and the Bcl-G BH3 domain (Figure S2B). We hypothesized that the BH3 domain of Bcl-G could also be sufficient to block Bak dependent apoptosis. tBID^Bcl-G^ and tBID^BNIP5^ were expressed lower than their inactive mutants (Figure S2C), however tBID^Bcl-G^ but not tBID^Bcl-G^ 3R was able to block apoptosis in Bax-deficient HeLa cells in response to BH3 mimetic treatment and promote long-term survival (Figure S2D-S2E).

To assess if expression of BNIP5 or tBID^BNIP5^ protects Bax-deficient HeLa from other apoptotic stimuli other than BH3 mimetics, we stably transduced Bax-deficient HeLa cells with constructs expressing either BNIP5 or tBID^BNIP5^, or with their inactive mutants and stimulated the cells with various apoptotic stimuli. Cell death was reduced in cells expressing BNIP5 or tBID^BNIP5^ but was unchanged in cells expressing ΔBH3 BNIP5 or tBID^BNIP5^-4R compared to parental cells (Figure 2G).

To evaluate whether BNIP5 expression results in enhanced tumor growth *in vivo*, we transduced CT.26 cells with BNIP5 or an empty vector and implanted these cells into Balb/c mice. Tumors from BNIP5-expressing cells grew larger compared to empty vector-expressing cells (Figure 2G-I). Together, these results indicate that the BH3 domain of BNIP5 is both necessary and sufficient to block Bak induced MOMP and apoptosis and promote tumor growth, and that the BH3 domain of Bcl-G is similarly sufficient to block Bak dependent apoptosis.

### Degradation of Bak upon BNIP5 expression is dispensable for inhibition of cell death

We noticed varying levels of Bak protein expression depending on the expression levels of BNIP5 in cell lines we generated. Higher levels of BNIP5 expression, driven by a retroviral promoter (pMX-BNIP5) in HeLa cells correlated with lower Bak protein levels, compared to cells expressing BNIP5 using the CRISPRa system or a CMV driven BNIP5 construct (Figure 3A). Lower Bak expression also correlated with stronger protection against BH3 mimetic-induced apoptosis (Figure 3B). Expression of tBID^BNIP5^ from the same promoter was sufficient to lead to degradation of endogenous Bak, while either ΔBH3 BNIP5 or tBID^BNIP5^-4R did not lead to degradation of Bak (Figure S3A). Similar results were obtained in HeLa cells expressing GFP-Bak and the constructs described above (Figure S3B), suggesting posttranscriptional regulation of Bak levels by BNIP5. We assessed if proteasomal degradation contributes to reduction of Bak levels in the presence of BNIP5. Treatment of BNIP5-expressing cells with either Bortezomib or MG132 increased Bak levels in a concentration-dependent manner (Figure 3C). However, treatment with these inhibitors did not sensitize BNIP5-expressing cells to induction of apoptosis upon co-treatment with BH3 mimetics (Figure 3D). To assess Bak function more directly under these conditions, we treated cells for 24 hours with proteasome inhibitors and measured cytochrome c release upon treatment with BH3 mimetics by flow cytometry. In alignment with previous results, cytochrome c was retained in mitochondria in BNIP5-expressing, Bax-deficient HeLa cells treated with proteasome inhibitors and BH3 mimetics, suggesting that Bak remains inhibited under these conditions (Figure 3E).

**Figure 3:**
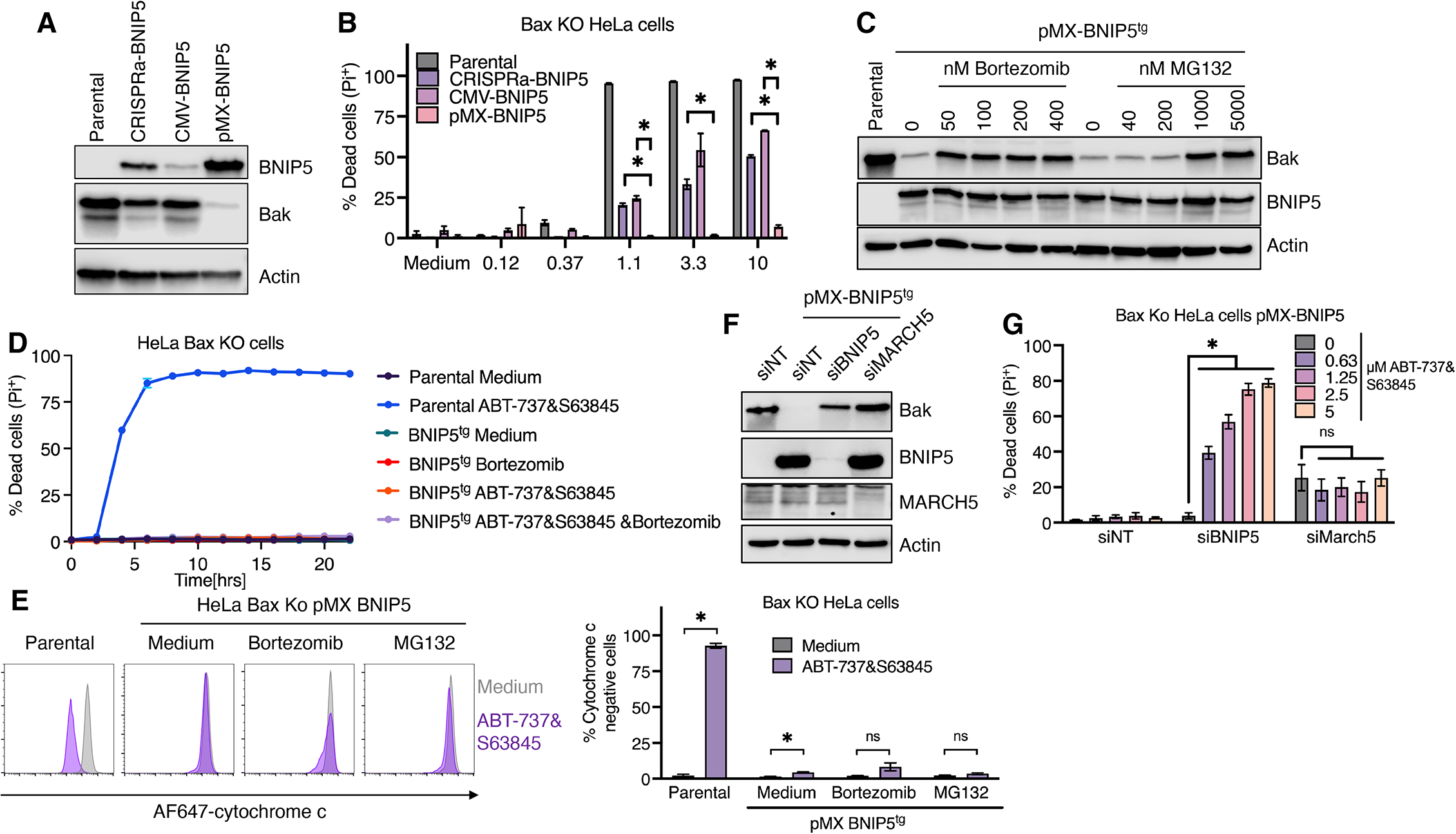
Degradation of Bak upon BNIP5 expression is dispensable for inhibition of cell death. **A** Western blot analysis of Bax deficient HeLa cells expressing various BNIP5 constructs. **B** Cells shown in **A** were treated with different concentrations of ABT-737 and S63845 and cell death was determined by measuring PI positive cells. **C** Western blot analysis of pMX-BNIP5 expressing, Bax deficient HeLa cells treated with indicated concentration of proteasome inhibitors for 24 hours. **D** PI incorporation of parental or pMX-BNIP5 expressing cells co-treated with BH3 mimetics and bortezomib. **E** Cytochrome c release from parental or pMX-BNIP5 expressing Bax deficient HeLa cells treated for 20 hours with indicated proteasome inhibitors and stimulated with BH3 mimetics for 4 hours. **F** & **G** pMX-BNIP5 expressing Bax deficient HeLa cells were transfected for 48 hours with indicated siRNAs and analyzed by western blot (**F**) or treated with increasing concentrations of BH3 mimetics and percentage of PI positive cells was enumerated **G**. * p<0.05. All panels show one experiment representative of at least 3 independent biological experiments.

The E3 ligase March5 has been shown to promote degradation of Bak-Mcl-1 complexes(Huang et al., 2022). To test if March5 regulates Bak degradation in the presence of BNIP5, we transfected pMX-BNIP5-expressing, Bax-deficient HeLa cells with siRNA targeting BNIP5 or March5. Expression of Bak increased upon transfection with either siRNA compared to a control siRNA (Figure 3F). Transfection of March5-targeting siRNA induced basal cell death in pMX-BNIP5-transduced, Bax-deficient HeLa cells, however only transfection with BNIP5 targeting siRNA restored sensitivity to BH3 mimetic induced apoptosis, indicating that Bak was still inhibited upon silencing of March5, despite basal cell death, in pMX-BNIP5-transduced Bax-deficient HeLa cells (Figure 3G). These results indicate that low BNIP5 expression is sufficient to effectively block Bak dependent apoptosis and that higher BNIP5 expression results in reduction of Bak protein levels, dependent on March5 and proteasomal degradation. However, this degradation seems to be dispensable for BNIP5-mediated Bak inhibition.

### The proteasome mediates removal of Bak from mitochondria in the presence of BNIP5

Based our previous results showing that degradation of Bak in the presence of BNIP5 is dispensable for inhibition of apoptosis, we hypothesized that BNIP5 expression changes the localization of Bak. We analyzed GFP-Bak localization in control or CRISPRa BNIP5-expressing cells transduced with GFP-Bak using immunofluorescence analysis. We found that less GFP-Bak localized to mitochondria in BNIP5-expressing cells compared to control cells (Figure 4A). Tom20 and Bak colocalization was quantified using the Pearson’s correlation coefficient (PCC), a metric independent of image intensity to correct for lower Bak levels in BNIP5 expressing cells (as described in STAR methods). To determine if this effect was dependent on the BH3 domain of BNIP5, we transiently expressed wildtype and ΔBH3 BNIP5 in GFP-Bak-expressing HeLa cells. GFP-Bak lost mitochondrial localization in the presence of wildtype but not ΔBH3 BNIP5, suggesting that the interaction with Bak mediated by the BNIP5 BH3 domain is necessary for translocation of Bak from mitochondria (Figure S4A and S4B). To rule out an effect of the bulky GFP tag, we validated these results using cells expressing HA-tagged Bak, which were transfected with BNIP5 (Figure S4C and S4D).

**Figure 4:**
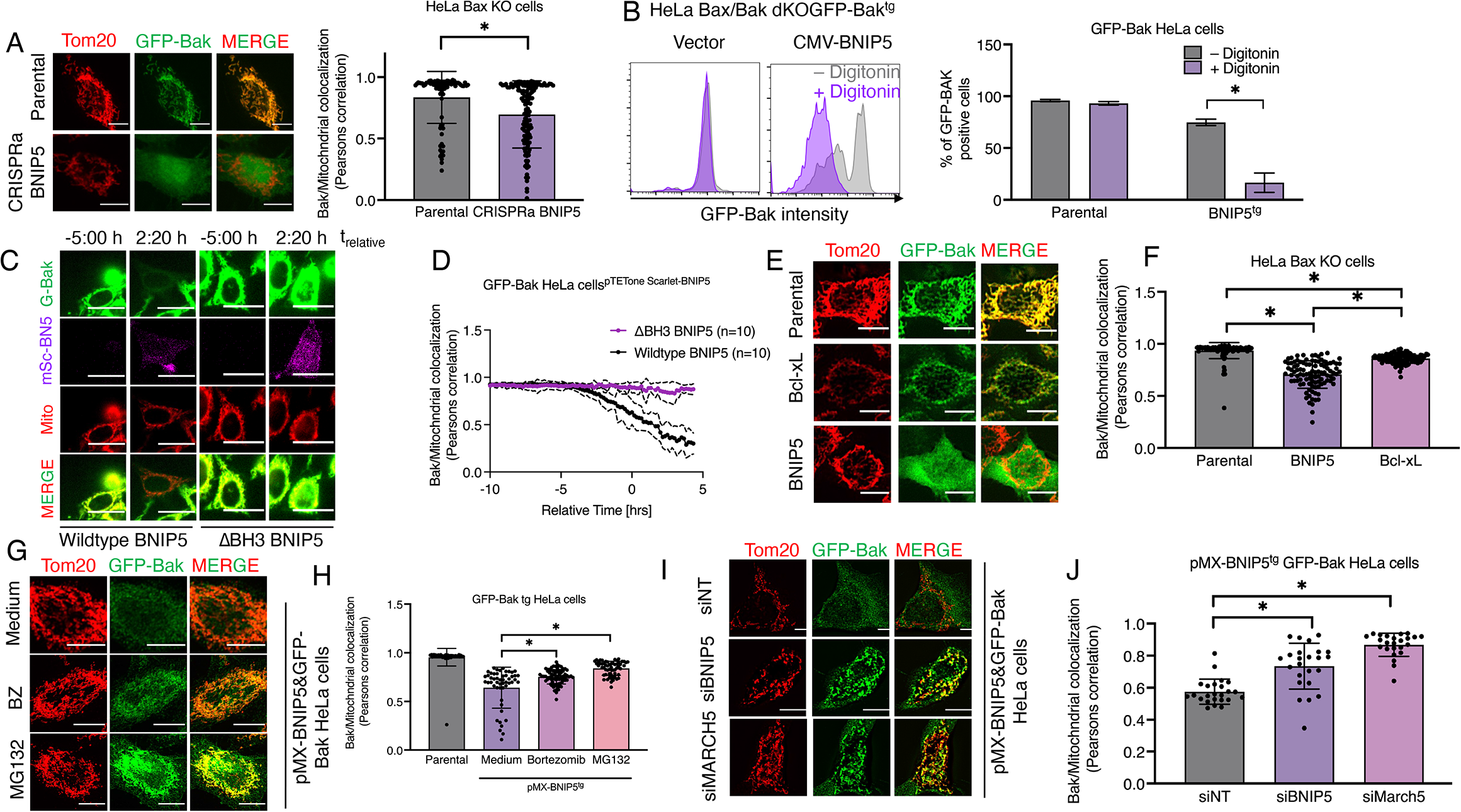
The proteasome mediates removal of Bak from mitochondria in the presence of BNIP5. **A** Immunofluorescence analysis of GFP-Bak and Tom20 in GFP-Bak or CRISPRa-BNIP5/GFP-Bak transgenic cells. Pearson’s correlation coefficient as a proxy of Bak and mitochondrial colocalization is quantified. **B** Flow Cytometry analysis (and quantification thereof) of Bax deficient HeLa cells stably expressing a control plasmid or a CMV driven human BNIP5 construct before (gray) or after (purple) permeabilization with digitonin. **C**&**D** Live cell microscopy of GFP-Bak, pTetONE^3G^ mScarlet-ΔBH3 or mScarlet-WT-BNIP5 expressing Bax/Bak double deficient HeLa cells. Time after doxycycline addition is shown and normalized to onset of BNIP5 expression. Ten representative cells of each BNIP5 genotype are quantified in **D**. Relative time was normalized to onset of BNIP5 expression. **E-F** Tom20 and GFP immunofluorescence of GFP-Bak transgenic or GFP-Bak and BNIP5/Bcl-xL expressing cells. Colocalization of Tom20 and GFP was quantified in at least 100 cells and is shown **F**. **G**-**H** GFP-Bak and Tom20 immunofluorescence analysis of indicated cells, either left untreated or treated with 1 µM Bortezomib or 5 µM MG132 for 6 hours. Quantification of at least 100 cells is shown in **H**. **I**-**J** GFP-Bak and Tom20 immunofluorescence analysis of pMX-BNIP5 expressing Bax deficient HeLa cells transfected with the indicated siRNAs for 48 hours. Quantification of 25 cells per condition is shown in **J**. All panels show one experiment representative of at least 3 independent biological experiments. Scale bars for **A, E, G, I** are 10 µm, and 30 µm for **C**.

To test if the BH3 domain of BNIP5 is sufficient to promote stable removal from mitochondria, we transfected tBID^BNIP5^ or tBID^BNIP5^-4R into GFP-Bak-expressing HeLa cells and measured colocalization of Bak and mitochondria. Expression of tBID^BNIP5^ reduced colocalization of Bak and mitochondria (Figure S4E and S4F). Stable expression of tBID^Bcl-G^ also led to stable removal of GFP-Bak from mitochondria (Figure S4G and S4H), suggesting a similar function of the BH3 domain of Bcl-G. To assess if GFP-Bak was integrated into the OMM or cytosolic, BNIP5- and GFP-Bak-expressing cells were permeabilized with a low concentration of digitonin to remove soluble GFP-Bak and subsequently analyzed by flow cytometry. In cells expressing GFP-Bak, most fluorescence was retained after permeabilization, however in cells co-expressing BNIP5, very little GFP-Bak was retained after treatment with digitonin, indicating that GFP-Bak was no longer inserted into membranes in the presence of BNIP5 (Figure 4B).

To assess the kinetics of Bak removal by BNIP5, we created doxycycline inducible mScarlet-BNIP5 fusion proteins and performed live-cell imaging in GFP-Bak-expressing cells exposed to doxycycline, where we labeled mitochondria using MitoTracker deep red. Cells expressing wildtype BNIP5 showed translocation of GFP-Bak from mitochondria within a period of 2-4 hours of detectable BNIP5 expression (Figure 4C, 4D and Movie S1), while in cells expressing ΔBH3 BNIP5, GFP-Bak remained localized to mitochondria (Figure 4C, 4D and Movie S2).

Multidomain Bcl-2 proteins are shuttled between the OMM and the cytosol in a process called retrotranslocation, and Bcl-xL has been shown to cause retrotranslocation of Bak to and from the OMM (Edlich et al., 2011; Todt et al., 2015). However, cells stably expressing Bcl-xL showed a significantly more mitochondrial distribution of GFP-Bak compared to cells expressing BNIP5 (Figure 4F and G) suggesting that BNIP5 more permanently removes Bak from mitochondria.

Alternatively, BNIP5 could mediate removal of Bak from mitochondria through blocking translocation of newly synthesized Bak molecules to mitochondria. If this is the mechanism of action of BNIP5, blocking translation of Bak in the absence of BNIP5 expression should be sufficient to result in stable removal of Bak from mitochondria, as ‘old’ Bak molecules are constitutively removed from mitochondria and synthesis of new molecules is blocked. We treated pTET-One3G Scarlet-BNIP5 and GFP-Bak transduced cells with either doxycycline (Dox) or cycloheximide (Chx) for 16 hours and imaged GFP-Bak localization. Only expression of BNIP5 through addition of doxycycline lead to reduced colocalization of GFP-Bak and mitotracker deep red staining. Blocking Bak synthesis by Chx treatment in the absence of BNIP5 induction did not change GFP-Bak localization at similar time points, when expression of BNIP5 removed Bak from mitochondria (Figure S4I and S4J, Movies S3-S5). These results suggest that de novo GFP-BAK synthesis contributes minimally to its mitochondrial levels whereas BNIP5 leads to stable removal of Bak from mitochondria rather than enhanced retrotranslocation or inhibition of localization of only newly formed Bak molecules.

The proteasome has been proposed to cooperate with other co-factors in extraction of proteins from membranes(Basch et al., 2020; Karbowski and Youle, 2011) Treatment of pMX-BNIP5, GFP-Bak-expressing HeLa cells with either bortezomib or MG132 for only 6 hours promoted GFP-Bak localization to mitochondria (Figure 4G and H). Depletion of either BNIP5 or MARCH5 in the same cells also increased colocalization of Bak and mitochondria (Figure 4I and J). Treatment with proteasome inhibitors or depletion of March5 did not reverse inhibition of Bak by BNIP5 (Figure 3 D-G) suggesting that cytosolic translocation by the proteasome of Bak is dispensable for BNIP5 mediated inhibition of apoptosis.

### BNIP5 activates Bak to engage MODE 2 inhibition

We tested if expression of BNIP5 is sufficient to block Bak activation in the absence of known Bcl-2 family members. HCT116 allBcl2KO cells are devoid of 17 known Bcl-2 family proteins (O’Neill et al., 2016). Ectopic expression of Bak or Bax in these cells results in auto-activation of the effector proteins and MOMP (O’Neill et al., 2016). We generated HCT116 allBcl-2KO cells expressing Bcl-xL or BNIP5. Subsequent transduction of cells with retrovirus encoding either GFP, GFP-Bak or GFP-Bax revealed that BNIP5 expression does not increase the fraction of viable, GFP-positive cells compared to parental HCT116 allBcl-2KO cells. Bcl-xL-expressing cells displayed 10-times more viable GFP-Bax or GFP-Bak positive cells (Figure 5A and S5A). This suggest that BNIP5 requires other Bcl-2 family proteins to inhibit Bak activation.

**Figure 5:**
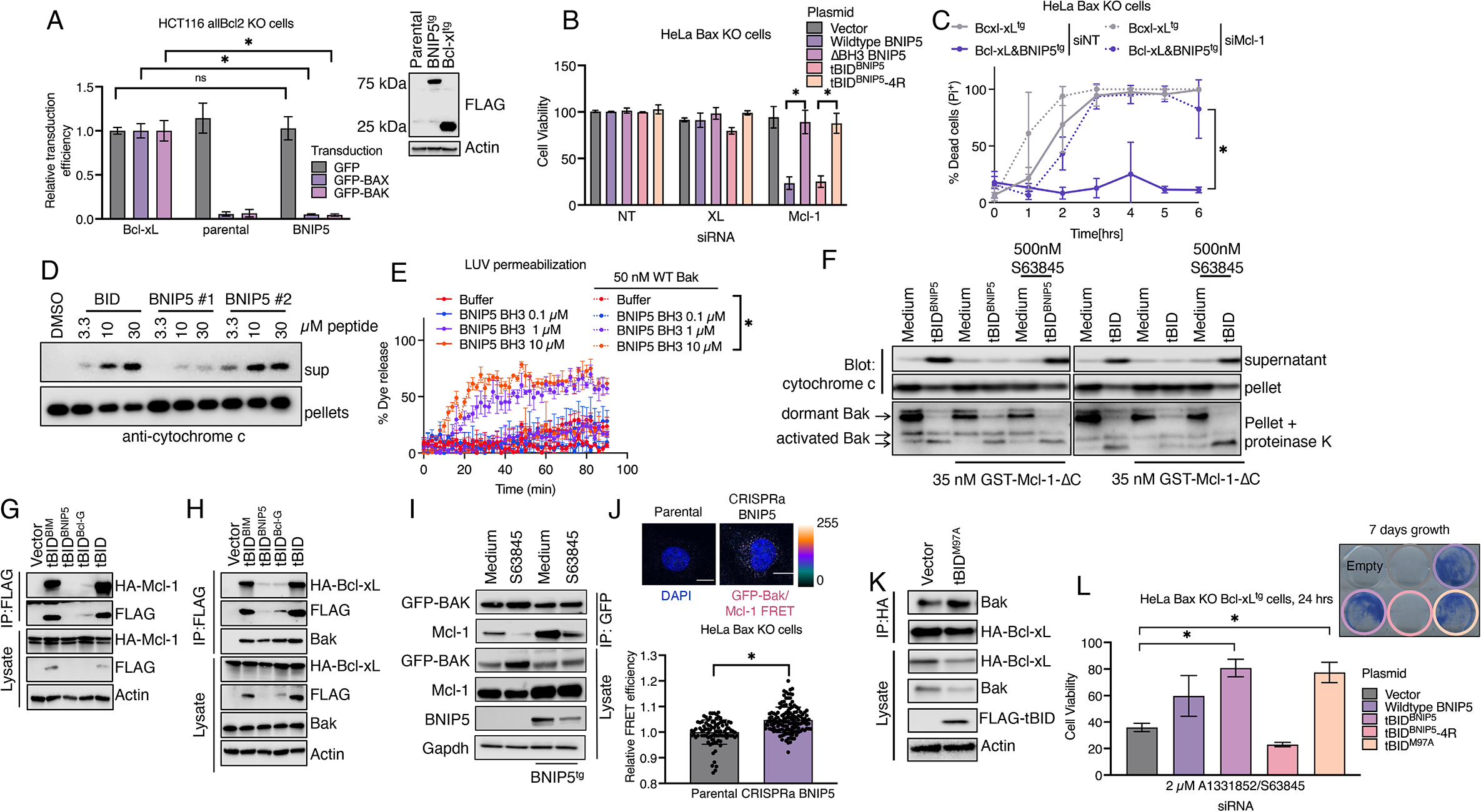
Anti-apoptotic BH3 only proteins activate Bak to drive MODE 2 inhibition. **A** Parental, Bcl-xL- or BNIP5-expressing HCT116 allBcl-2 KO cells were transduced with GFP, GFP-Bak or GFP-Bax encoding retrovirus and number of live, GFP-positive cells was enumerated using flow cytometry (for representative raw data see Figure **S3A**). **B** ATP-based cell viability assay of Bax-deficient HeLa cells at 24 hours after co-transfection of indicated siRNAs (X-axis) and plasmids (colors). **C** Cell death as measured by number of PI positive cells of indicated cell lines transfected with either control or Mcl-1 targeting siRNA and stimulated with 2 µM A-1331852 and S63845. **D** Cytochrome C release from mitochondrial heavy membrane fractions purified from HCT-116 cells was assessed 90 minutes after treatment with peptides at indicated concentrations. **E** Dye release from LUVs incubated with recombinant Bak and the indicated peptides. **F** Limited proteinase k proteolysis and cytochrome c release from MLMs treated with IVTT tBID chimeras. **G-H** Immunoprecipitation of FLAG-tBID chimeras from transiently transfected HEK 293T. Cells were co-transfected with HA-Mcl-1 (**G**) or HA-Bcl-xL (**H**). **I** Immunoprecipitation of GFP-Bak from Bax, Bak double-deficient HeLa cells or cells expressing a CMV promoter driven BNIP5 construct. **J** Intermolecular FRET efficiency (between Mcl-1 and GFP-Bak) of parental or CRISPRa-BNIP5, Bax-deficient HeLa cells expressing GFP-Bak. **K** Immunoprecipitation of HA-Bcl-xL from transiently transfected HEK293T cells 48 hours post transfection. **L** ATP-based cell viability (24 hours post stimulation) and methylene blue staining (7 days post stimulation) of Bax-deficient, Bcl-xL-expressing HeLa cells transfected with the indicated constructs for 24 hours and treated with 5 µM ABT-737&S63845. **A, B, C, J** show pooled data of three independent biological replicates. All other panels show one experiment representative of at least three independent experiments. * p<0.05

Mcl-1 and Bcl-xL are the major anti-apoptotic Bcl-2 proteins, which can inhibit Bak-dependent apoptosis (Kale et al., 2018). To investigate the role of Bcl-xL and Mcl-1 in BNIP5-mediated Bak inhibition, we co-transfected BNIP5, tBID^BNIP5^ or their inactive mutants with siRNA targeting Mcl-1 or Bcl-xL into Bax-deficient HeLa cells. Simultaneous expression of BNIP5 and knock-down of Mcl-1 or Bcl-xL revealed that expression of BNIP5 or tBID^BNIP5^ in the absence of Mcl-1, but not Bcl-xL, led to spontaneous loss of cell viability in Bax-deficient HeLa cells. Expression of ΔBH3 BNIP5 or tBID^BNIP5^-4R had no effect on cell viability (Figure 5B). Knock-down efficiency of Mcl-1 and Bcl-xL was not affected by transfection of any plasmid (Figure S5B).

To allow for more efficient Bcl-xL depletion over a longer time, we repeated these results using a doxycycline inducible BNIP5 construct. Bax-deficient HeLa cells were transfected for 48 hours with siRNA against Mcl-1, Bcl-xL or co-transfected with siRNA against Bak and treated for 24 hours with doxycycline. While silencing of Bcl-xL or Mcl-1 had no effect on cell viability in the absence of doxycycline, silencing of Mcl-1 but not Bcl-xL resulted in loss of cell viability upon induction of BNIP5 by doxycycline treatment. This loss in cell viability was prevented by co-depletion of Bak (Figure S5C and S5D). Expression of Bcl-xL and Mcl-1, but not an inactive Mcl-1 mutant harboring mutations in the BH3 groove (A209R, L213R, V216R, V220R) prevented loss of cell-viability induced by BNIP5 in the absence of endogenous Mcl-1 (Figure S5E). These results indicate that expression of BNIP5 in the absence of Mcl-1, or sufficient levels of Bcl-xL, leads to activation of Bak and apoptosis.

To interrogate if complexes formed by Bak and Bcl-xL in the presence of BNIP5 resist BH3 mimetic treatment, we generated Bcl-xL- and BNIP5-expressing cells and tested BH3 mimetic sensitivity in cells transfected with control siRNA or siRNA targeting Mcl-1. BNIP5-expressing cells overexpressing Bcl-xL were not protected from BH3 mimetic-induced apoptosis upon depletion of Mcl-1 expression by siRNA, suggesting that Mcl-1 is required for BNIP5-mediated Bak inhibition even in the presence of excess Bcl-xL (Figure 5C and Figure S5F). These results suggest that BNIP5 expression induces Bak-dependent apoptosis in the absence of Mcl-1 or Bcl-xL and that only complexes formed by Bak and Mcl-1 protect cells from BH3 mimetic-induced apoptosis.

We interrogated the ability of the BNIP5 BH3 domain to modulate Bak activation in purified mitochondria. These mitochondrial preparations are devoid of cytosolic Bax (Uren et al., 2007). Treatment of mitochondria with two BH3 peptides derived from BNIP5 or BID indicated that all peptides can induce cytochrome c release in a concentration dependent manner (Figure 5D), suggesting that the BNIP5 BH3 functions as a direct activator of Bak, rather than an inhibitor. We then assessed the ability of the BNIP5 BH3 to activate recombinant Bak using dye release from large unilamelar vesicles (LUVs) (Singh et al., 2022). Treatment of LUVs containing dormant Bak with BNIP5 peptides induced concentration dependent dye release, suggestion that the BNIP5 BH3 domain activates recombinant Bak (Figure 5E). To investigate if BNIP5-induced Bak activation can be blocked by Mcl-1, we treated purified mitochondria with tBID^BNIP5^ or tBID in the presence of purified Mcl-1-ΔC. Mcl-1 blocked cytochrome c release induced by either tBID or tBID^BNIP5^, while tBID^BNIP5^-4R did not induce cytochrome c release (Figure S5G). To test if tBID^BNIP5^ can activate Bak in the presence of Mcl-1, we subjected mitochondria treated with tBID^BNIP5^ or tBID to a limited proteolysis assay using proteinase k (Hockings et al., 2018). Protease sensitivity of Bak was used as a measure to assess if Bak had transitioned to an ‘open’ stated. Our antibody cannot distinguish fully active Bak or ‘open’ Bak bound to Mcl-1(Moldoveanu et al., 2013). Bak was processed by proteinase k in the presence of tBID^BNIP5^ and Mcl-1, although cytochrome c release was blocked. Pre-incubation with low concentrations of S63845 restored cytochrome c release (Figure 5F). Similar results were obtained with tBID^Bcl-G^ (Figure S5H). These results suggest that the BH3 domains of BNIP5 and Bcl-G activate Bak and promote an open, proteinase k sensitive conformation and engage MODE 2 inhibition *in vitro*.

Based on the results discussed above we tested if BH3 domains of BNIP5 and Bcl-G interact with Bak to activate it, while not binding to Mcl-1. To test this hypothesis in cells, we transiently co-expressed FLAG tagged tBID chimeras with HA-Mcl-1 or HA-Bcl-xL in HEK293T cells. Immunoprecipitation of tBID and tBID^BIM^ confirmed strong interaction between these constructs and Mcl-1, while tBID^BNIP5^ and tBID^Bcl-G^ failed to bind Mcl-1. However, tBID^BNIP5^ and tBID^Bcl-G^ were expressed at lower levels than tBID and tBID^BIM^ (Figure 5G). Similar results were obtained if the interaction with HA-Bcl-xL was analyzed by immunoprecipitation. We concomitantly tested the ability of all constructs to interact with Bak and found that comparable amounts of Bak were immunoprecipitated by all tBID chimeras (Figure 5H). This suggests that, although expressed at lower levels, tBID^BNIP5^ and tBID^Bcl-G^ can potently interact with Bak while sparing anti-apoptotic proteins, therefore identifying them as highly potent Bak-activating BH3 domains.

If BNIP5 acts to activate Bak, but apoptosis is subsequently blocked, we hypothesized that more Bak will be found in MODE 2 inhibition i.e., bound to Mcl-1. Immunoprecipitation experiments from Bax-deficient HeLa cells expressing GFP-Bak and a CMV promoter driven BNIP5 construct, allowing for Bak inhibition with moderate degradation of Bak, revealed that more Mcl-1 precipitated with GFP-Bak in the presence of BNIP5. Treatment with BH3 mimetics showed that also under these conditions, more Mcl-1 remained bound to GFP-Bak in the presence of BNIP5 despite the presence of the Mcl-1 inhibitor (Figure 5I). Transient transfection of HEK293T cells with HA-Bak, BNIP5 and FLAG-Mcl-1 showed that more Bak precipitated with Mcl-1 in the presence of BNIP5, both in the absence and presence of S636845 (Figure S5I).

We then used a Förster resonance energy transfer (FRET) based assay to examine Bak-Mcl-1 interaction in fixed cells. Cells expressing GFP-Bak or GFP-Bak and BNIP5 were stained for GFP and endogenous Mcl-1 using Alexa Fluor 488- and Alexa Fluor 555-conjugated secondary antibodies respectively. FRET transfer from Alexa Fluor 488 to Alexa Fluor 555 was measured to determine the relative amount of GFP-Bak bound to Mcl-1 in the presence and absence of BNIP5. Intermolecular FRET efficiency was higher in the presence of BNIP5 compared to parental cells (Figure 5J) confirming enhanced binding of Bak to Mcl-1 in the presence of BNIP5.

Engagement of MODE 2 inhibition has been proposed to make cells more resistant to apoptosis induction, as complexes formed between anti-apoptotic proteins and activated effectors are more stable than MODE 1 complexes between BH3-only proteins and anti-apoptotic proteins (Hockings et al., 2018). We used a tBID mutant (tBID^M97A^), reported to have a low affinity for anti-apoptotic proteins but capable of activating Bak (Lee et al., 2016). Co-expression of tBID^M97A^ and Bcl-xL in HEK293T cells increased the interaction of Bak and Bcl-xL (Figure 5K). tBID^M97A^ was cytotoxic in HeLa cells (data not shown), so we used Bcl-xL-expressing, Bax-deficient HeLa cells to study its impact on MODE 2 engagement in cells. Bcl-xL-expressing HeLa cells transfected with tBID^M97A^ showed increased cell viability (24 hours) and clonogenic survival (7 days) in response to BH3 mimetic treatment, to an extent comparable to transfection of different BNIP5 constructs (Figure 5L).

Overall, our results indicate that ectopic expression of Bcl-G and BNIP5 potently inhibits Bak dependent apoptosis in various cells. Removal of Bak from mitochondria as well as proteasomal Bak degradation was dispensable for the function of BNIP5. However, BNIP5 and Bcl-G harbor BH3 domains, which act as selective Bak activators, while not inhibiting anti-apoptotic proteins. This leads to increased formation of MODE 2 complexes between Bak and Mcl-1, which are resistant to disruption by apoptotic stimuli (Figure S6).

## Discussion

Our results describe the existence of uncharacterized BH3-only proteins which can inhibit Bak dependent apoptosis through engagement of MODE 2 inhibition. Several studies have suggested that if Bax or Bak are bound to anti-apoptotic proteins, i.e. in MODE 2 inhibition, cells are more resistant to apoptosis induction compared to cells where the effectors are dormant and anti-apoptotic proteins are bound to proapoptotic BH3-only proteins, i.e. in MODE 1 inhibition(Hockings et al., 2018; Llambi et al., 2011). The discovery of the BH3 domains of BNIP5 and Bcl-G demonstrates that naturally occurring proteins can induce MODE 2 inhibition in cells to promote apoptosis resistance.

Our results also suggest that BH3 mimetics need to act on anti-apoptotic proteins that are either unoccupied by BH3-only proteins or transiently bound to the latter. On purified mitochondria, only high (> 30 µM) ABT-737 concentrations were able to dislodge Bcl-xL-Bak complexes (Hockings et al., 2018). We found that pretreatment at nanomolar concentrations with BH3 mimetics is sufficient to induce MOMP upon Bak activation (Figure 5F). This suggests that in cells, a small fraction of Bak molecules is activated in resting conditions, which needs to be neutralized by anti-apoptotic proteins. Addition of BH3 mimetics to cells occupies free anti-apoptotic proteins, preventing neutralization of active Bak (or Bax) molecules, which leads to apoptosis initiation. Therefore, enhanced binding of Bak to Mcl-1 in the presence of anti-apoptotic BH3-only proteins does not allow BH3 mimetics (or other apoptotic stimuli) to readily induce MOMP and apoptosis.

Since all the experiments presented in this study rely on overexpression of proteins, we are unable to determine if our results extend to the function of endogenous BNIP5. Unfortunately, were unable to find cell lines which expressed detectable protein levels of BNIP5 (data not shown). Publicly available gene expression data show that BNIP5 is expressed selectively along tissues of the gastrointestinal tract and that expression strongly correlates with Bak mRNA expression (Yue et al., 2014). This suggests that at least both proteins are present in the same tissues and therefore regulation of Bak by BNIP5 is conceivable under these circumstances.

Different models of the interactions between Bcl-2 proteins have been proposed to date(Chen et al., 2015; Huang et al., 2019; Llambi et al., 2011; O’Neill et al., 2016) but our understanding of how the core apoptotic machinery interacts with other survival pathways remains limited. Multiple influences outside the Bcl-2 family, including various metabolic pathways(Pollyea et al., 2018; Stevens et al., 2020), Hexokinase II (Lauterwasser et al., 2021; Majewski et al., 2004b, 2004a; Schindler and Foley, 2013) mitochondrial cristae integrity (Chen et al., 2019) or DRP1 (Jenner et al., 2022) have been suggested to regulate MOMP. One could also imagine that transcriptional repressors might exist that prevent the expression of Bax and Bak. Our genome-wide transactivation CRISPR-dCas9 screen, along with a complementary knock-out CRISPR-Cas9 screen (Chin et al., 2018) revealed that survival in response to engagement of the core apoptotic machinery by BH3 mimetics is almost exclusively dependent on Bcl-2 proteins. We identified only Bcl2A1, an anti-apoptotic protein not antagonized by our BH3 mimetics prevented apoptosis in cells expressing Bax. The fact that our screening effort failed to recover such regulators suggests that the magnitude of apoptosis regulation ascribed to most pathways outside the Bcl-2 family might be minimal during direct engagement of high levels of MOMP by BH3 mimetics. The concentrations and stimulation times used in our screening approach were comparably high, potentially masking more subtle regulatory pathways. This however again suggests that survival of cells and ultimately tumor relapse or progression are regulated by the Bcl-2 family in response to BH3 mimetics and to a large degree in response to chemotherapy (Chonghaile et al., 2011).

Bax and Bak have been viewed as largely functionally redundant molecules. The expression of Bax being required for male reproduction in mice (Knudson et al., 1995), to our knowledge is the only demonstration of one effector having an exclusive function *in vivo*. Why then, do all vertebrate animals have Bak? The BH3 domains of BNIP5 and Bcl-G identified in this study can selectively activate Bak, drive MODE 2 inhibition via Bak-Mcl-1 interaction and hence inhibit Bak dependent apoptosis. VDAC2 has been identified by a CRISPR-Cas9 knock-out screen to be required for Bax dependent apoptosis (Chin et al., 2018). These results show that only regulation of a single effector can be achieved by such mechanisms. Microbes that evade apoptosis to replicate can do so via inhibition of Bax and Bak, but mechanisms that target only one effector (e.g., the inhibition of Bax by the herpesvirus protein VMIA (Arnoult et al., 2004) will be relatively ineffective when both effectors are present. Our study suggests that such general inhibition can only be effectively mediated by expression of anti-apoptotic Bcl-2 proteins (as observed for the viral Bcl-2 homologs(Fitzsimmons et al., 2020; Kvansakul et al., 2007; Loh et al., 2005). This may provide a molecular explanation as to why two MOMP effector proteins have been conserved in these animals.

Our results indicate that that unidentified Bcl-2 family members exist which can function in regulation of intrinsic apoptosis through unconventional inhibition of Bak-dependent apoptosis. The mechanisms described here also suggests that direct activation of MOMP effectors is not sufficient to trigger apoptosis but that the concomitant inhibition of anti-apoptotic proteins is required for Bak oligomerization and activation. Bak activation by BNIP5 or Bcl-G, both containing BH3 domains which do not neutralize Mcl-1, instead inhibits subsequent Bak activation through increased binding of Bak to Mcl-1. This explains why potent inducers of apoptosis such as Bim or tBid can directly activate Bak and de-repress Bak-Mcl-1 interactions. Pro-apoptotic BH3-only sensitizer proteins such as Bad or Noxa hence represent a crucial component of the Bcl-2 family for efficient induction of apoptosis.

## Acknowledgements

We thank Xiaofei Wang and Tanya Khan for the help with animal experiments. We thank Dr. Katherine Verbist for proofreading the manuscript.

## Author Contributions

S.R and D.R.G conceived the study, S.R. performed most experiments, T.M., G.C. and Z.L. performed experiments. S.R. and D.R.G wrote the initial draft of the manuscript. All authors commented on the manuscript. D.R.G. supervised the research.

## Methods

### Cell lines and culture conditions

HeLa, A375, HCT116 and HEK293T cells were obtained from ATCC and maintained in DMEM 10 %FCS, 2 mM L-Glutamine, 1% Penicillin-Streptomycin at 37 °C and 5% CO_2._ PC9 and CT.26 cells were maintained in RPMI 10 % Fetal Calf Serum (FCS), 2m L-Glutamine, 1% Penicillin-Streptomycin. Cells were subcultured every 3-4 days, authenticated in house and regularly tested for mycoplasma contamination using Mycoalert kit (Lonza).

### Cell Lysis and Western Blot analysis

At the end of the experiment, cells were washed once using cold PBS and lysed in RIPA buffer (300 mM NaCL, 1mM EDTA, 1% NP-40, 0.5% Sodium deoxycholate, 0.1% SDS, 50 mM Tris-HCl pH 7.4) supplemented with HALT protease and phosphatase inhibitor cocktail (Thermo Scientific) and lysed for 30 minutes on ice. Lysates were clarified through centrifugation at 21’000 *xg* at 4 °C and supernatants were transferred to a new tube. Protein concentration was measured by BCA assay (Thermo Scientific) and samples were adjusted to the lowest concentration. 30-50 µg of protein per lane were separated on CRITERION 4-12% SDS polyacrylamide gels (Biorad) and transferred to 0.2 µm nitrocellulose membrane (GE Healthcare) using semi dry transfer. Membranes were blocked for 1 hour in 3% milk in TBST and incubated with primary antibodies overnight at 4 °C. After three washes with TBST, membranes were incubated with HRP conjugated secondary antibodies for 1 hour at room temperature. Membranes were washed again three times with TBST and developed using chemiluminescent substrate (ECL-Clarity, Biorad or ECL-Sirius, Advansta).

### Cell viability and cell death measurements

One day prior to the experiment, cells were seeded at 5×10^3^ cells per well of a flat-bottom 96-well plate in 100 µl of medium and allowed to adhere overnight. For cell death measurements, cells were labeled with Syto16 and Sytox Orange (Thermo Scientific) and stimulated with indicated stimuli. Images of cells were acquired using an Incucyte system (Essen Biosciences) and percentage of dead (Sytox orange +) cells relative to all cells (Syto16+) was calculated. For cell viability assays, CellTiterGlo (Promega) assay was diluted 1:4 in PBS and 100 µl were added to each well of a 96-well plate and processed according to the manufacturer’s instructions. For long-term survival assays, 5×10^3^ cells were seeded per well of a 24-well plate and stimulated as described above. Stimuli were removed (after times indicated in the figure legends) and cells were cultured for 7-10 days until untreated controls reached confluency. Cells were stained with 0.5% Methylene blue in 50% methanol/PBS for 30 minutes at room temperature and washed 3 times with DI water prior to acquisition of pictures.

### CRISPR-Cas9 transactivation screen

HeLa cells were transduced with a dCas9-VP64 encoding plasmid and selected with 5 µg/ml Blasticidin for 7 days. Blasticidin concentration was reduced to 1 µg/ml. Human Calabrese CRISPRa p65-HSF library was obtained from addgene and amplified. Lentiviral particles were produced in HEK293T cells and after 48 hours, viral supernatant was harvested and frozen to −80°C. Viral titers were determined as described in Ref. Cells were infected at an MOI of 0.25 and after 48 hours they were selected using 2 µg/ml Puromycin for 7-10 days. 5×10^7^ cells were harvested as a pre-stimulation control. Cells were subjected to 3 rounds of stimulation with 20xIC_50_ of ABT-737&S63845 for 16 hours. Cells were allowed to recover and stimulation was repeated twice more. Outgrowing cells were harvested and genomic DNA was isolated using the Qiagen Blood&Tissue MaxiPrep. SgRNAs sequences were amplified by PCR, purified by gel-extraction and analyzed by next-generation sequencing. Reads were aligned using pynaplpy (ref) and normalized counts were analyzed using custom python scripts.

#### Cytochrome c release assay measured by flow cytometry

HeLa cells were seeded one day prior to the experiment at 7.5 x10^4^ cells per well of a 24-well plate in 500 µl growth medium. The next day, cells were treated with drugs in the presence of 20 µM QVD-OPh to prevent caspase activation. After 4-6 hours, cells were harvested by trypsinization and transferred to 5 ml round bottom polystyrene tubes. Cells were then washed once with cold PBS and permeabilized with 500 µl of 25 µg/ml digitonin in PBS for 10 minutes on ice. Cells were fixed using BD fix and perm buffer for 15 minutes at 4 °C. After 2 washed with BD wash and perm buffer, cells were stained with 50 µl of Alexa-Fluor 647 conjugated anti-Cytochrome-C antibody (1:500, BD) at room temperature for 30 minutes. Cells were washed again two times with BD wash and perm buffer and analyzed on an Cytek Aurora spectral flow cytometer.

#### Cytochrome c release from mitochondrial heavy membrane fractions

Mitochondrial heavy membrane fractions were isolated from HCT116 (REF) cells or murine livers (Ref) as described elsewhere. Purified mitochondria were resuspended in AT buffer (300 mM trehalose, 10 mM HEPES–KOH pH 7.7, 10 mM KCl, 1 mM EDTA and 0.2% BSA, 90 mM KCl) concentration and 100 µg/50 µl reaction were incubated with peptides or tBID chimeras for 90 minutes at 37 °C. Pellet and supernatant was separated by centrifugation (5000 *xg*, 10 minutes, 4 °C) and supernatants were mixed with 4x western blot sample buffer. Pellets were washed again and resuspended in 1x western blot sample buffer. Samples were boiled at 95 °C for 5 minutes and analyzed by SDS Page as described above.

### Generation of knock-out cell lines using CRISPR-Cas9

sgRNAS (Table 1) were cloned into pX458(addgene) and transfected into HeLa or MEF cells as described above. 24-48 hours post-transfection, transfected cells were sorted by FACS based on GFP expression. Cells were allowed to recove for 7 days after sorting and either used as pools or single clones were generated by limiting dilution into 96-well plates. Deletion was confirmed by western blot.

### Generation of plasmids for mammalian expression and amplification

All ORFs were either synthesized by genscript or amplified from cDNA libraries. PCR products were cloned into either pLJM1 (for lentiviral expression), pcDNA 3.1 (for transient expression) or pMX-IRES-Hygromycin/Puromycin (for retroviral expression) using Takara-Clontech in-fusion cloning kit. Assembled plasmids were transformed into Stellar chemically competent *E.Coli* (Takara-Clontech) and plated on appropriate Luria Bertani agar plates. Correct clones were verified by sequencing and plasmids were amplified using Zymo Research endotoxin free midi prep kit.

### Generation of stable cell lines

5x 10^6^ HEK 293T cells were seeded per 10 cm tissue culture treated dish the day before the experiment. The next day, cells were transfected with 9 µg packaging plasmids (pCL Ampho for retroviral particles or psPAX2 for lentiviral particles), 9 µg transfer vector and 2 µg pVSV-G using 60 µl linear PEI (Polysciences). Medium was exchanged 6-8 hours post-transfection and harvested after another 48 hours. Viral supernatant was mixed with polybrene (8 µg/ml final concentration) and 1x 10^6^ cells were infected using 10 ml of lentiviral supernatant or spin-infected (1900 *xg*, 45 minutes) with 10 ml of retroviral supernatant. Cells were selected with appropriate antibiotic or sorted by FACS 2-3 days post transduction.

### Immunofluorescence and colocalization analysis

On the day prior to the experiment, poly-L-lysine coated glass coverslips (#1.5, EMS) were placed into tissue culture treated 12-well plates and washed twice with growth medium and cells were seeded at 1.5 x 10^5^ per well. At the end of the experiment, cells were fixed with 4% Paraformaldehyde (PFA, EMS) for 15 minutes at room temperature. PFA was quenched and cells were permeabilized with 100 mM Glycine, 0.1% Triton in PBS for 5 minutes. After 3 washes with PBS, cells were blocked in PBS, 5% FCS for 30 minutes. Coverslips were stained in a humid chamber with primary antibodies diluted in PBS, 5% FCS overnight at 4 °C. Coverslips were washed three times with PBS and stained with Alexa-Fluor-Plus conjugated secondary antibodies (Thermo Scientific) for 1 hour at room temperature. Coverslips were washed again as before and mounted using anti-fade glass mounting medium supplemented with NucBlue (Thermo Scientitic). Mounted slides were imaged using a Marianas spinning disk confocal (Intelligent Imaging Innovations) comprised of an inverted AxioObserver Z.1 (Zeiss), CSU-W with SoRA (Yokogawa), Prime95B sCMOS camera (Photometrics), and solid-state laser illumination with wavelengths as appropriate. Cells were segmented based on mitochondrial staining using Cellpose(Stringer et al., 2021) and Pearson’s correlation coefficient (PCC) was calculated for every plane along the z-axis between the Tom20 and Bak channel, for every cell using the formula

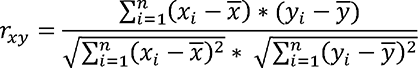

where

x = the linearized image vector for channel 1

y = the linearized image vector for channel 2

n = the length of both vectors

*x̅* = the mean of x

*y̅* = the mean of y

An average of all PCCs was calculated and used as PCC per cells. All these calculations were performed using custom scripts which are available upon request.

### Immunoprecipitation in HEK293T cells

1×106 HEK293T cells per well of a 6-well plate were seeded the day before the experiment (3 wells per condition). The next day cells were transfected with a total amount of 3 µg of plasmid DNA using 10 µl of Lipofectamine 2000 according to the manufacturer’s instructions. 48 hours post-transfection, cells were washed once with cold PBS and lysed in IP buffer (20 mM Tris-HCl pH 7.4, 142.5 mM KCl, 2 mM CaCl2, 1% CHAPS, Protease inhibitors) for 20 minutes on ice. Lysates were clarified by centrifugation (21’000 xg at 4 °C) and equal amounts of supernatant were transferred to a new tube and 5% of lysate was removed as input control. 20 µl of FLAG-M2 magnetic beads (Sigma Aldrich) or HA-magnetic beads (Pierce) per reaction were washed once with IP buffer and added to lysates. Reactions were incubated at 4 °C for 4 hours on an end-over-end rotator. Beads were washed 4 times 5 minutes with IP buffer using end-over-end rotation and proteins were eluted by resuspension of beads in PBS supplemented with 1% SDS. Beads were boiled for 3 minutes at 95 °C, eluates were mixed with 4x sample buffer and boiled again. Lysates and immunoprecipitations were separated by SDS and analyzed as described above.

### Liposome permeabilization assays

Large unilamellar vesicle (LUVs) or liposomes were prepared using lipid films of the following lipid composition resembling that of mitochondrial outer membrane: 26.6% phosphatidylethanolamine, 40.9% phosphatidylcholine, 8.3% phosphatidylserine, 9.1% phosphatidylinositol, 7% cardiolipin and 8.0% Ni2+-affinity lipid 1,2-dioleoyl-sn-glycero-3-[(N-(5-amino-1-carboxypentyl)iminodiacetic acid)succinyl] (nickel salt) [DGS NTA(Ni), 2.5 μM final concentration in each assay] (Avanti Polar Lipids). The fluorescent dye 12.5 mM ANTS (8-aminonaphthalene-1, 3, 6-trisulfonic acid, disodium salt and and quencher 45 mM DPX (p-xylene-bis-pyridiniumbromide), were encapsulated in liposomes by extrusion of the resuspended film in LUV buffer (10 mM HEPES pH 6.8, 5 mM MgCl_2_, 200 mM KCl) containing ANTS/DPX. LUVs were separated from excess ANTS/DPX by S500 size exclusion chromatography and were stored at 4 °C in the dark. Assays were setup in 96-well black flat bottom plates (Costar) on ice and LUV permeabilization was monitored at 37 °C for 90 min in a CLARIOSTAR microplate reader (BMG Labtech) at excitation and emission wavelengths of 360 nm and 530 nm, respectively. Data were normalized relative to minimum and maximum fluorescence induced by buffer and 5% CHAPS, respectively.

### Live cell imaging

GFP-BAK expressing, TetOne3G BNIP5 transduced HeLa cells were seeded at 75’000 cells per well of a 24-well ibidi chamber and grown overnight in full medium at 37 °C, 5% CO_2_. Mitochondria were labelled with 50 nM Mitotracker Deep Red for 30 minutes at 37 °C, 5% CO_2_ in full medium. Cells were washed once with phenol red free DMEM and doxycycline (2 µg/ml) or cycloheximide (1 µg/ml) in phenol red free DMEM, 10% FBS, 1% Hepes were added. Cells were imaged every 10 minutes for 16 hours. Bak

### Recombinant protein expression and purification

Mcl-1ΔTM was cloned into pGEX4T1 and transformed into NEB T7 express *E.Coli* (NEB). One clone was inoculated overnight in 200 ml LB+100 µg/ml ampicillin. The next day, 8 l of LB+ampicillin were inoculated with 160 ml overnight culture and grown at 37 °C until an OD_600_ of 0.8. Protein expression was induced using 1 mM IPTG overnight at 20 °C. Bacteria were harvested by centrifugation and resuspended in 100 ml 300 mM NaCl, 25 mM Tris pH 7.5, 1 mM PMSF and lysed by sonication. Lysates were clarified by centrifugation (15’000 *xg*, 4 °C, 30 minutes) and supernatants were incubated with 1 ml Glutathione Sepharose (GE Healthcare) for 2 hours at 4 °C. Beads were washed 3 times 5 minutes using 50 ml wash buffer (150 mM NaCl, 25 mM Tris pH 7.5) by end-over-end rotation. Proteins were eluted using 5 mM reduced glutathione in wash buffer by incubating beads 3 times 10 minutes on an end-over-end rotator at room temperature. GST-Mcl-1ΔTM was further purified using size exclusion chromatography. Pooled fractions were concentrated and flash frozen to be stored at −80 °C in wash buffer supplemented with 10% glycerol.

### siRNA and plasmid transfections

Cells were seeded at 20% confluency in 6-well plates one day prior to transfection. The next day, cells were transfected using 50 pmols siRNA and 9 µl of Lipofectamine RNAimax (Thermo Scientific) according to the manufacturer’s instructions. 48-72 hours post-transfection, cells were lifted by trypsinization counted and seeded for respective assays. For plasmid transfections, cells were seeded to a confluency of 80-90% one day prior to transfection and transfected the next day with a total of 2.5 µg of endotoxin-free plasmid DNA using 7.5-9 µl of Lipofectamine 2000 according to the manufacturer’s instructions. 24 hours post-transfection, cells were lifted, counted, and seeded for cell viability assays and western blot analysis. For immunofluorescence analysis cells were seeded (1.5 x 10^5^ HeLa cells) directly on poly-L-lysine coated coverslips (Electron Microscopy Sciences), which were placed in a 12-well plate. Cells were transfected using 1 µg of DNA and 2 µl of Lipofectamine 2000 according to the manufacturer’s instructions.

### Tumor studies

5x 10^5^ CT.26 cells suspended in PBS were implanted into the dorsal flank of female, 8-week old Balb/c mice, purchased from Jackson Laboratories (Maine, US). Tumor growth was monitored using a digital caliber every 3-4 days by measuring the short (s) and long axis (l) of the tumor. Tumor volume was calculated as l x s^2^. At Day 25 post implantation, mice were sacrificed by CO2 inhalation tumors were isolated, weighed and pictures were taken. All mice were bred and housed in specific pathogen-free facilities, in a 12-hour light/dark cycle in ventilated cages, with chow and water supply ad libitum, at the Animal Resources Center at St. Jude Children’s Research Hospital. Mouse studies were conducted in accordance with protocols approved by the St. Jude Children’s Research Hospital Committee on Care and Use of Animals and in compliance with all relevant ethical guidelines.

### Supplemental Figure Legends

**Supplemental Figure 1:**

**A** Western Blot analysis of HeLa dCas9-VP64 cells expressing indicated sgRNAs. PC9 (**B-D**) or A375 (**E-G)** dCas9-VP64 cells expressing sgRNAs targeting BNIP5 or Bcl-G were transfected for 72 hours with indicated siRNAs and stimulated with ABT-737 and S63845. Relative cell viability to untreated controls is shown in **B** and **E.** Western blot of BNIP5 expressing cells (**C** and **F**) or Bcl-G (**D** and **G**) transgenic cells are shown. **H** Bax-deficient MEFs expressing human or murine BNIP5 were stimulated with 5 µM ABT-737 and S63845 and dead cells were quantified using and Incucyte live cell imaging system. Cells were also analyzed for FLAG-BNIP5 expression. * p<0.05 **B, E, H** show pooled data of three independent biological experiments. All other panels show one experiment representative of at least two independent biological experiments.

**Supplemental Figure 2:**

**A** Western blot analysis of expression of different BNIP5 truncation mutants. Numbers in **B** indicate the amino acids of BNIP5 present in the respective mutant **B** Alignment of BNIP5 and Bcl-G BH3 domains. **C** Expression of FLAG-tBID chimeras was analyzed by western blot. **D** Bax-deficient HeLa cells were transfected with indicated tBID chimeras and stimulated with 2 µM ABT-737 and S63845 for 24 hours. Relative cell viability to untreated samples is shown. **E** Cells were treated as in **D** but survival was measured after 7 days using methylene blue staining. **D** shows pooled data of three independent experiments. **B, E, F** show data representative of at least two independent experiments.

**Supplemental Figure 3:**

Western blot analysis of Bax-deficient (**A**) of Bax/Bak double deficient GFP-Bak expressing (**B**) HeLa cells transduced with indicated constructs (expressed from a retroviral promoter). All results are representative at least 2 independent experiments.

**Supplemental Figure 4:**

**A**-**F** GFP-Bak transgenic (**A, B, E, F**) or HA-Bak (**C**, **D**) transgenic HeLa Bax/Bak double deficient cells were transfected with the indicated plasmid and colocalization of Bak with mitochondrial Tom20 staining was analyzed by immunofluorescence analysis. Quantification of each condition is shown in **B**, **D** and **F**. **G**-**H** GFP-Bak and Tom20 immunofluorescence analysis of GFP-Bak or GFP-Bak and tBID^Bcl-G^ expressing HeLa cells. **I-J** Live cell microscopy of GFP-Bak, pTetONE^3G^ WT-BNIP5 transgenic cells left either untreated, treated with doxycycline or cycloheximide for 16 hours. Colocalization of GFP-Bak and mitotracker deep red was quantified. Time was normalized to onset of BNIP5 expression. At least 50 cells per condition, across 3 independent experiments were quantified for **B** to **H**. Scale bars for **A-G** are 10 µm and 20 µm for **I** * p<0.05. Inset in **E** shows intensity matched GFP-Bak image.

**Supplemental Figure 5:**

**A** One experiment of data shown in Figure 5A. Parental, Bcl-xL or BNIP5 transgenic HCT116 allBcl-2 KO cells were transduced with GFP, GFP-Bak or GFP-Bax encoding retrovirus. Transduction efficiency (GFP, x-axis) and dead cells (DAPI, y-axis) are shown for one representative sample per condition. **B** Bax-deficient HeLa cells were co-transfected with indicated constructs and siRNAs for 24 hours in the presence of 40 µM qVD (to prevent cell death). Lysates were analyzed by western blot for indicated proteins. **C** and **D** HeLa Bax KO pTETone3G BNIP5 cells were transfected with siRNAs for 48 hours and subsequently treated for 16 hours with 2 µg/ml doxycycline. ATP-based cell viability (**C**) or expression analysis by western blot (**D**) was performed. **E** ATP based cell viability assay of Bax-deficient HeLa cells co-transfected with indicated siRNAs (x-axis) and combinations of plasmids (colors). **F** Expression analysis of Bax deficient HeLa cells expressing Bcl-xL or Bcl-xL and BNIP5 transfected with Mcl-1 targeting siRNA after 48 hours. **G** Cytochrome c release from murine liver mitochondria (MLMS, 50 µg per reaction) treated with indicated IVTT tBID chimeras for 90 minutes. **H** MLMs (300 µg per reaction) were treated with the indicated tBID chimeras (1% vol/vol), purified GST-Mcl-1ΔC and/or S63845 as described in main text. Cytochrome c release and Bak conformation (by limited proteolysis) was assayed after 90 minutes. **I** FLAG-Mcl-1 immunoprecipitation from transiently transfected HEK293T cells. * p<0.05. **C** and **E** show pooled data from 3 independent experiments. All other panels show one experiment representative of at least 3 independent biological experiments.

**Supplemental Figure 6**

**A** Model of the function of anti-apoptotic BH3-only proteins. In wildtype cells, Bak is found dormant on mitochondria. Apoptosis is engaged in response to BH3 mimetics as pro-apoptotic BH3-only proteins are displaced from anti-apoptotic proteins and hence the BH groove of anti-apoptotic proteins is occupied by BH3 mimetics (Top). In cells expressing anti-apoptotic BH3 only proteins, more Bak is bound to Mcl-1, a complex resistant to displacement by BH3 mimetics or other apoptotic stimuli. These complexes are removed by the proteasome from the OMM and Bak is degraded (Bottom).

**Movie S1**

Live cell imaging of Bax deficient HeLa, pTET3G-BNIP5(WT) and GFP-Bak(Green) expressing cells treated with doxycycline at start of the movie. Mitochondria were labelled with Mitotracker deep red (Cyan). Time is shown as hh:mm:ss. Scale bar is 10 µm.

**Movie S2**

Live cell imaging of Bax deficient HeLa, pTET3G-BNIP5(ΔBH3) and GFP-Bak(Green) expressing cells treated with doxycycline at start of the movie. Mitochondria were labelled with Mitotracker deep red (Cyan). Time is shown as hh:mm:ss. Scale bar is 10 µm.

**Movie S3**

Live cell imaging of Bax deficient HeLa, pTET3G-BNIP5(WT) and GFP-Bak(Green) expressing cells in medium. Mitochondria were labelled with Mitotracker deep red (Cyan). Time is shown as hh:mm:ss. Scale bar is 10 µm.

**Movie S4**

Live cell imaging of Bax deficient HeLa, pTET3G-BNIP5(WT) and GFP-Bak(Green) expressing cells treated with doxycycline at the start of the movie. Mitochondria were labelled with Mitotracker deep red (Cyan). Time is shown as hh:mm:ss. Scale bar is 10 µm.

**Movie S5**

Live cell imaging of Bax deficient HeLa, pTET3G-BNIP5(WT) and GFP-Bak(Green) expressing cells treated with cycloheximide at the start of the movie. Mitochondria were labelled with Mitotracker deep red (Cyan). Time is shown as hh:mm:ss. Scale bar is 10 µm.

