## Supplemental Figures and Legends for "Anti-apoptotic BH3-only proteins inhibit Bak-dependent apoptosis"

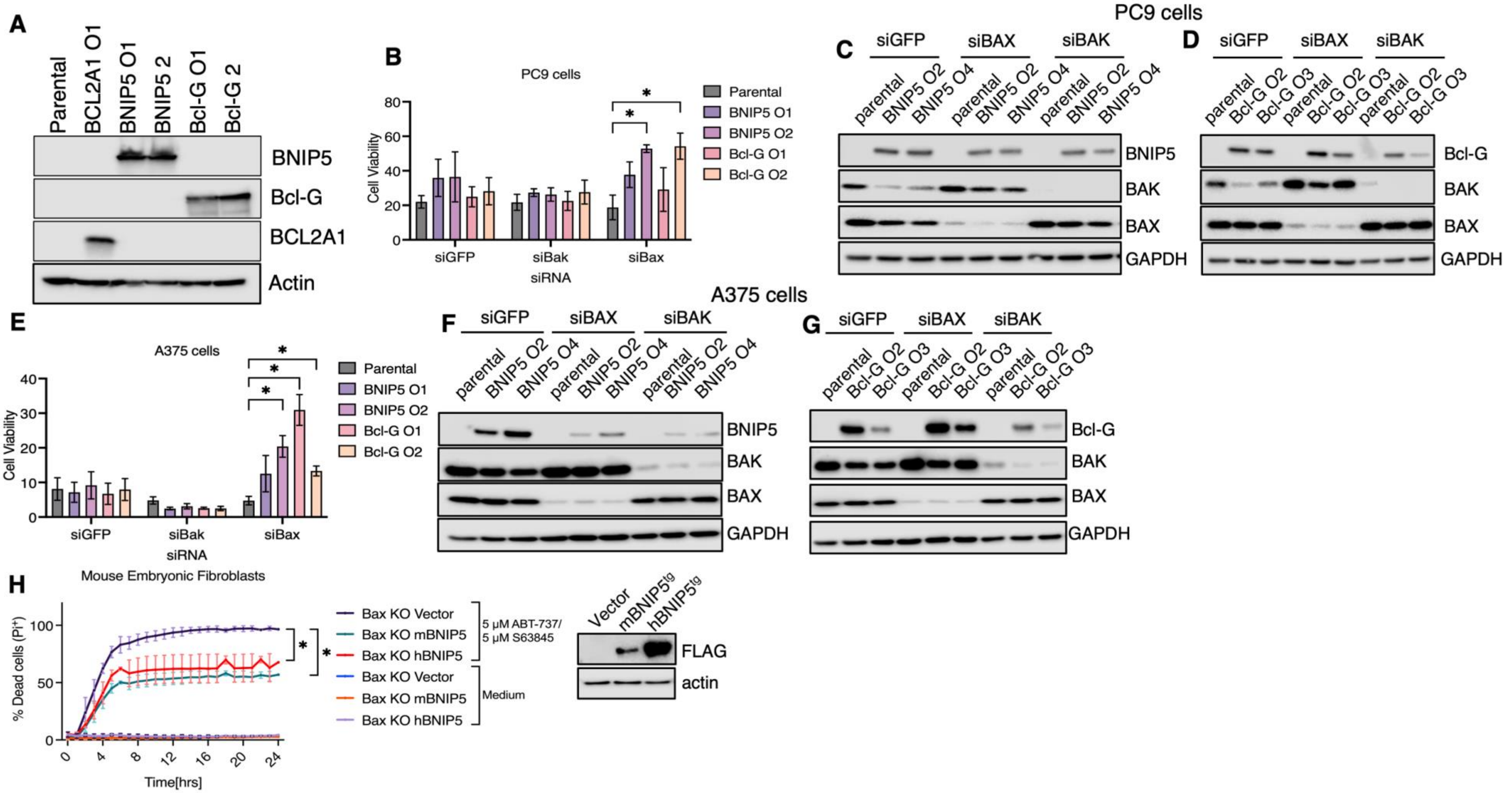


**Supplemental Figure 1:**

**A** Western Blot analysis of HeLa dCas9-VP64 cells expressing indicated sgRNAs. PC9 (**B-D**) or A375 (**E-G)** dCas9-VP64 cells expressing sgRNAs targeting BNIP5 or Bcl-G were transfected for 72 hours with indicated siRNAs and stimulated with ABT-737 and S63845. Relative cell viability to untreated controls is shown in **B** and **E.** Western blot of BNIP5 expressing cells (**C** and **F**) or Bcl-G (**D** and **G**) transgenic cells are shown. **H** Bax-deficient MEFs expressing human or murine BNIP5 were stimulated with 5 µM ABT-737 and S63845 and dead cells were quantified using and Incucyte live cell imaging system. Cells were also analyzed for FLAG-BNIP5 expression. * $p<0.05$ **B, E, H** show pooled data of three independent biological experiments. All other panels show one experiment representative of at least two independent biological experiments.

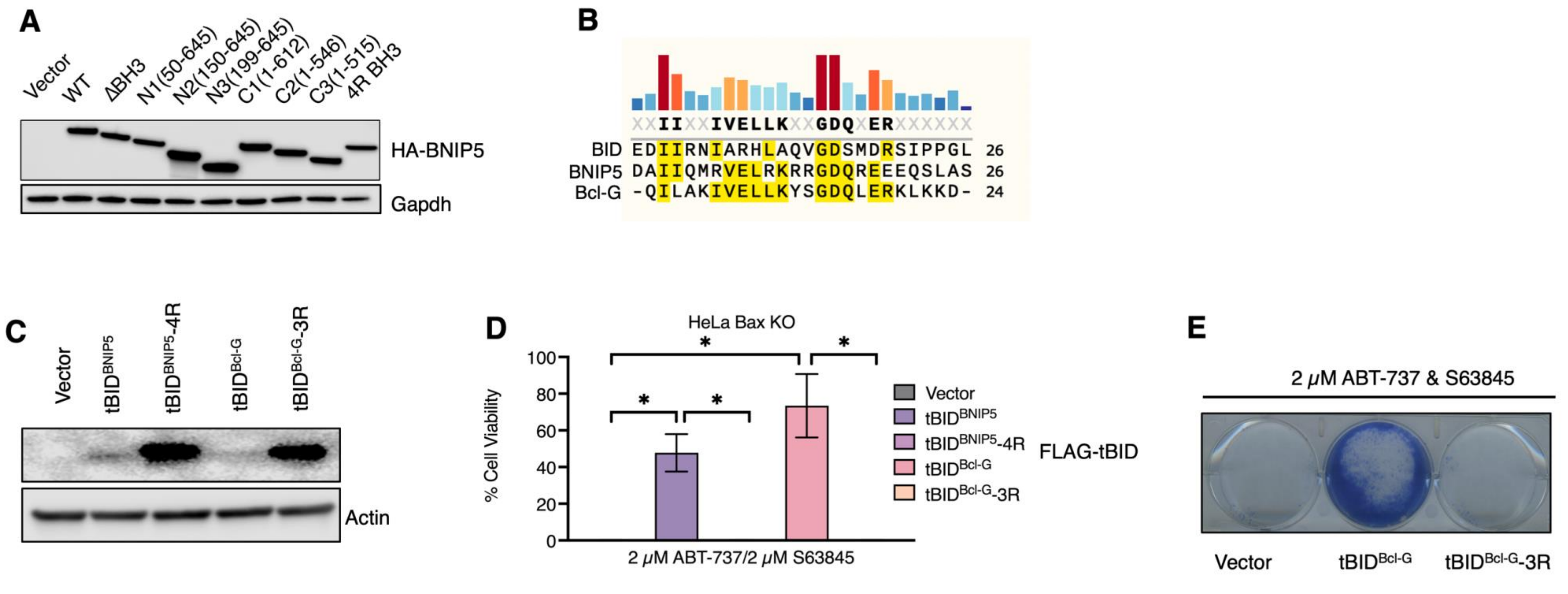


**Supplemental Figure 2:**
**A** Western blot analysis of expression of different BNIP5 truncation mutants. Numbers in **B** indicate the amino acids of BNIP5 present in the respective mutant **B** Alignment of BNIP5 and Bcl-G BH3 domains. **C** Expression of FLAG-tBID chimeras was analyzed by western blot. **D** Bax-deficient HeLa cells were transfected with indicated tBID chimeras and stimulated with 2 μM ABT-737 and S63845 for 24 hours. Relative cell viability to untreated samples is shown. **E** Cells were treated as in **D** but survival was measured after 7 days using methylene blue staining. **D** shows pooled data of three independent experiments. **B, E, F** show data representative of at least two independent experiments.

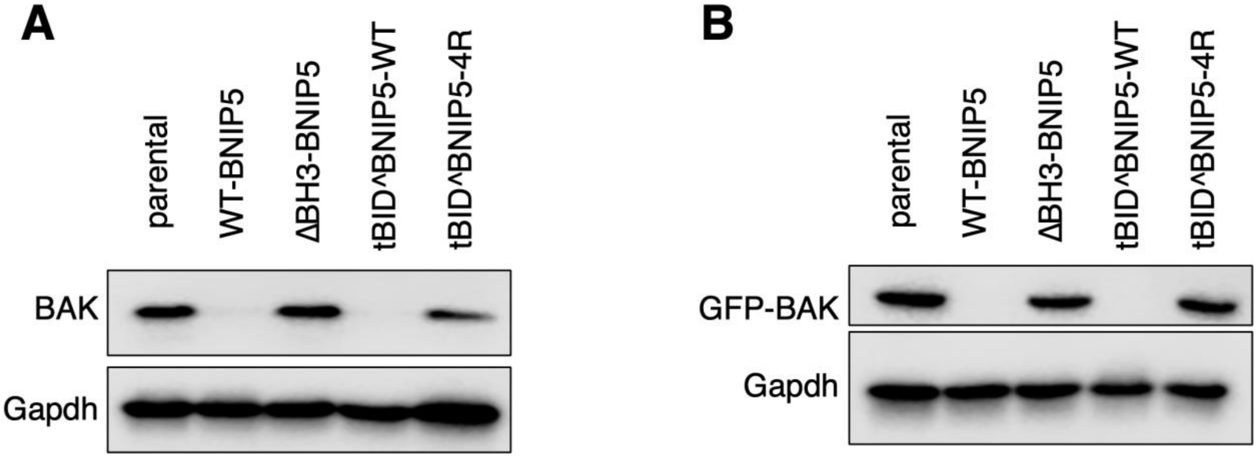


**Supplemental Figure 3:**
Western blot analysis of Bax-deficient (**A**) of Bax/Bak double deficient GFP-Bak expressing (**B**) HeLa cells transduced with indicated constructs (expressed from a retroviral promoter). All results are representative at least 2 independent experiments.

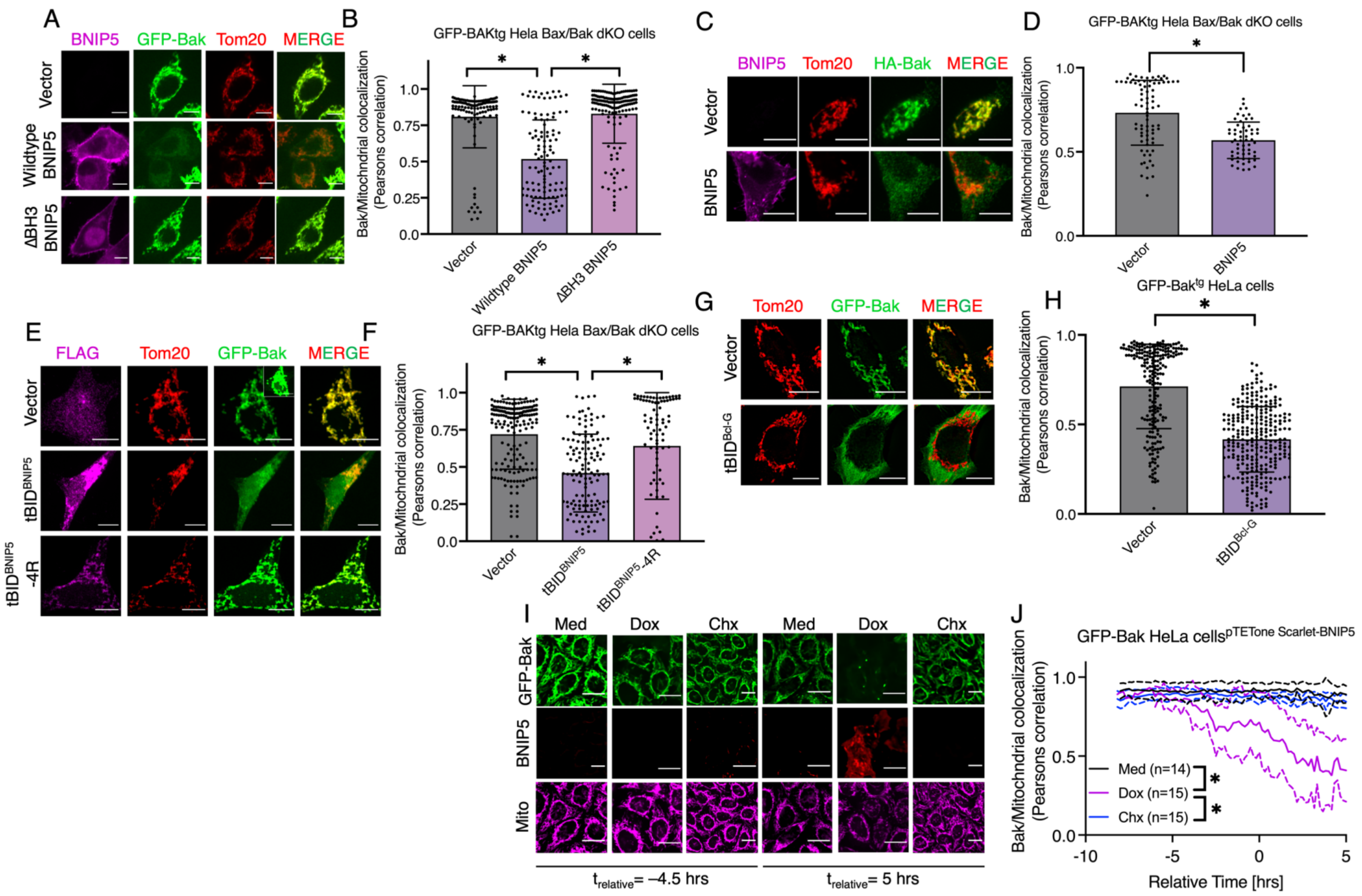


**Supplemental Figure 4:**

**A-F** GFP-Bak transgenic (**A, B, E, F**) or HA-Bak (**C, D**) transgenic HeLa Bax/Bak double deficient cells were transfected with the indicated plasmid and colocalization of Bak with mitochondrial Tom20 staining was analyzed by immunofluorescence analysis. Quantification of each condition is shown in **B**, **D** and **F**. **G-H** GFP-Bak and Tom20 immunofluorescence analysis of GFP-Bak or GFP-Bak and tBID$^{Bcl\text{-}G}$ expressing HeLa cells. **I-J** Live cell microscopy of GFP-Bak, pTetONE$^{3G}$ WT-BNIP5 transgenic cells left either untreated, treated with doxycycline or cycloheximide for 16 hours. Colocalization of GFP-Bak and mitotracker deep red was quantified. Time was normalized to onset of BNIP5 expression. At least 50 cells per condition, across 3 independent experiments were quantified for **B** to **H**. Scale bars for **A-G** are 10 µm and 20 µm for **I** * $p<0.05$. Inset in **E** shows intensity matched GFP-Bak image.

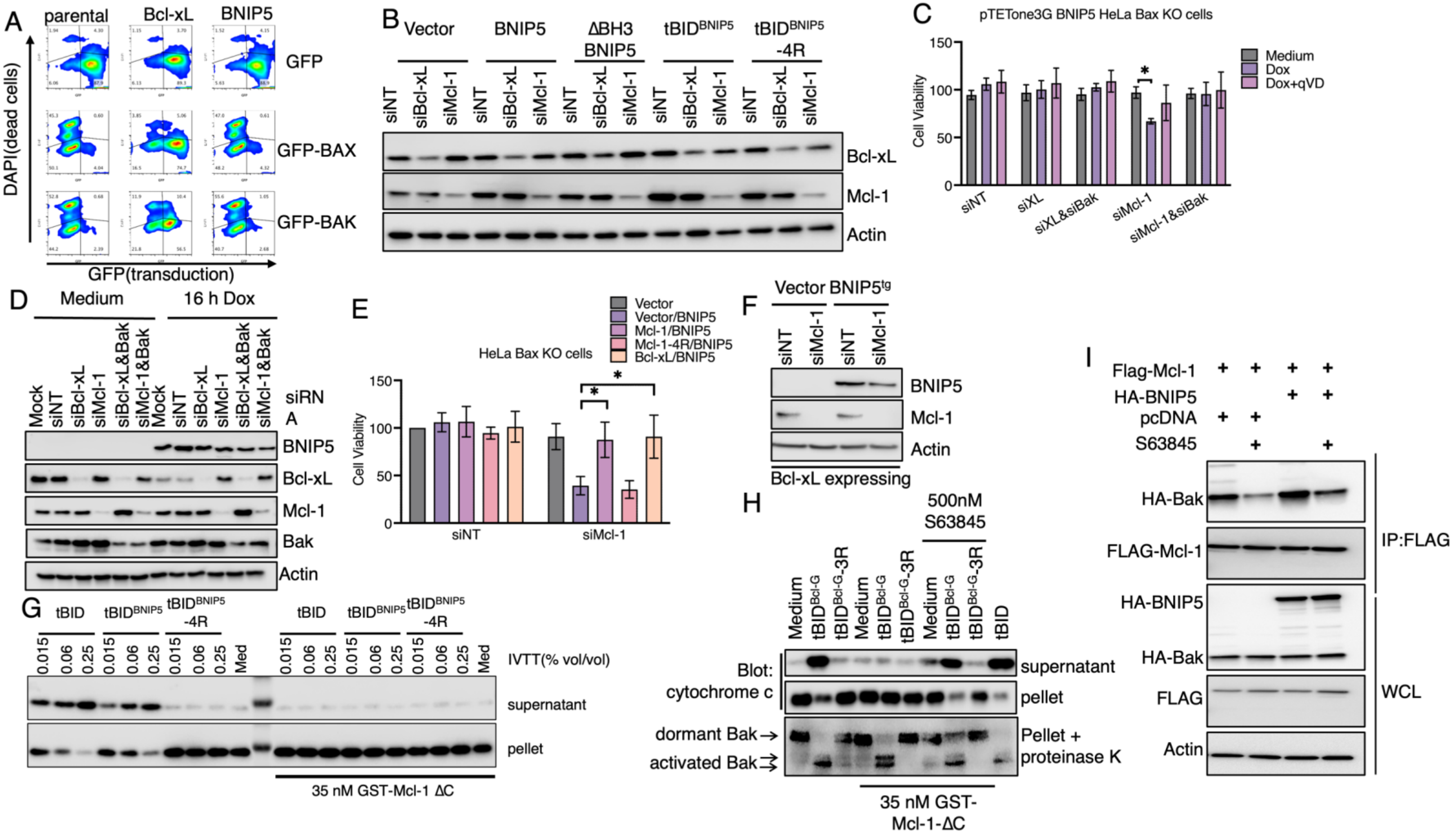

**Supplemental Figure 5:**

**A** One experiment of data shown in Figure **5A**. Parental, Bcl-xL or BNIP5 transgenic HCT116 allBcl-2 KO cells were transduced with GFP, GFP-Bak or GFP-Bax encoding retrovirus. Transduction efficiency (GFP, x-axis) and dead cells (DAPI, y-axis) are shown for one representative sample per condition. **B** Bax-deficient HeLa cells were co-transfected with indicated constructs and siRNAs for 24 hours in the presence of 40 µM qVD (to prevent cell death). Lysates were analyzed by western blot for indicated proteins. **C** and **D** HeLa Bax KO pTETone3G BNIP5 cells were transfected with siRNAs for 48 hours and subsequently treated for 16 hours with 2 µg/ml doxycycline. ATP-based cell viability (**C**) or expression analysis by western blot (**D**) was performed. **E** ATP based cell viability assay of Bax-deficient HeLa cells co-transfected with indicated siRNAs (x-axis) and combinations of plasmids (colors). **F** Expression analysis of Bax deficient HeLa cells expressing Bcl-xL or Bcl-xL and BNIP5 transfected with Mcl-1 targeting siRNA after 48 hours. **G** Cytochrome c release from murine liver mitochondria (MLMS, 50 µg per reaction) treated with indicated IVTT tBID chimeras for 90 minutes. **H** MLMs (300 µg per reaction) were treated with the indicated tBID chimeras (1% vol/vol), purified GST-Mcl-1ΔC and/or S63845 as described in main text. Cytochrome c release and Bak conformation (by limited proteolysis) was assayed after 90 minutes. **I** FLAG-Mcl-1 immunoprecipitation from transiently transfected HEK293T cells. * $p<0.05$. **C** and **E** show pooled data from 3 independent experiments. All other panels show one experiment representative of at least 3 independent biological experiments.

A

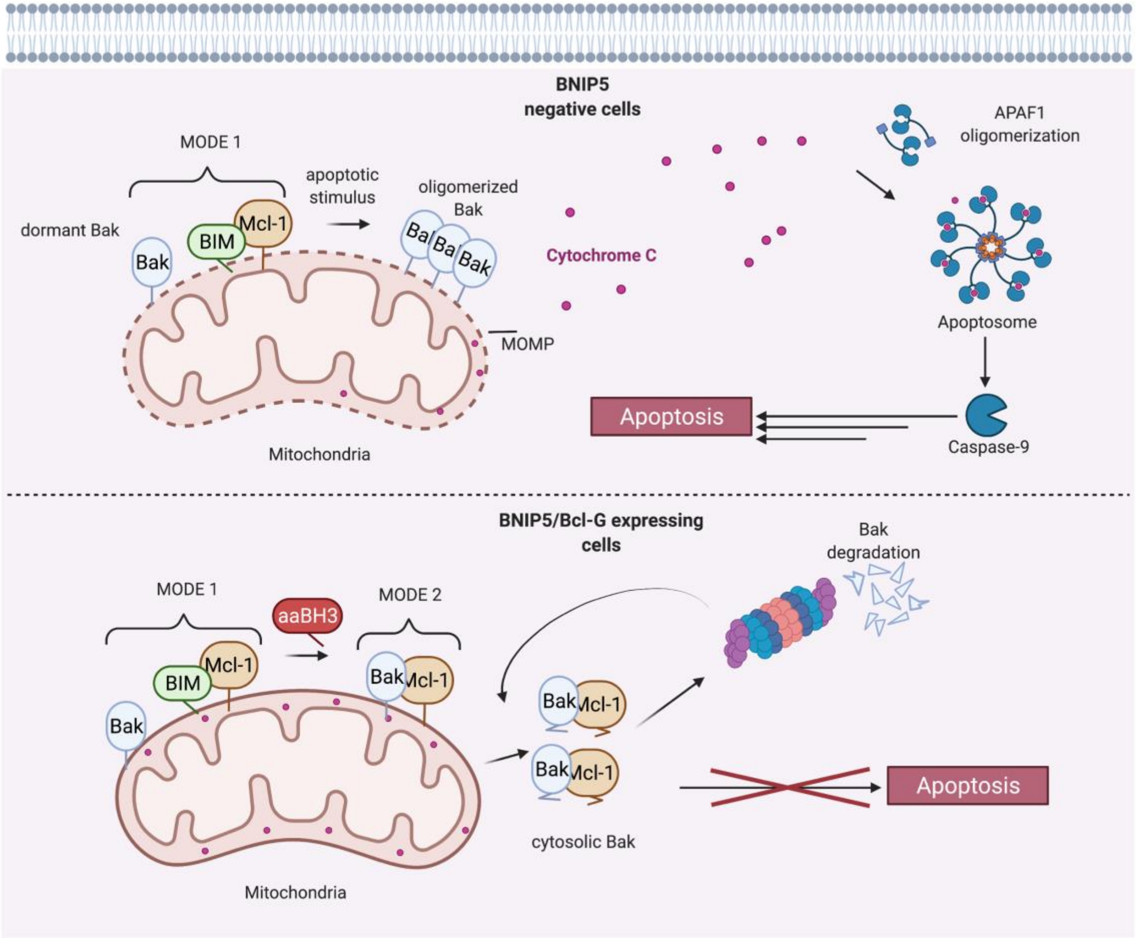


**Supplemental Figure 6**

**A** Model of the function of anti-apoptotic BH3-only proteins. In wildtype cells, Bak is found dormant on mitochondria. Apoptosis is engaged in response to BH3 mimetics as pro-apoptotic BH3-only proteins are displaced from anti-apoptotic proteins and hence the BH groove of anti-apoptotic proteins is occupied by BH3 mimetics (Top). In cells expressing anti-apoptotic BH3 only proteins, more Bak is bound to Mcl-1, a complex resistant to displacement by BH3 mimetics or other apoptotic stimuli. These complexes are removed by the proteasome from the OMM and Bak is degraded (Bottom).

**Supplemental Movie Legends:**

**Movie S1**

Live cell imaging of Bax deficient HeLa, pTET3G-BNIP5(WT) and GFP-Bak(Green) expressing cells treated with doxycycline at start of the movie. Mitochondria were labelled with Mitotracker deep red (Cyan). Time is shown as hh:mm:ss. Scale bar is 10 µm.

**Movie S2**

Live cell imaging of Bax deficient HeLa, pTET3G-BNIP5(ΔBH3) and GFP-Bak(Green) expressing cells treated with doxycycline at start of the movie. Mitochondria were labelled with Mitotracker deep red (Cyan). Time is shown as hh:mm:ss. Scale bar is 10 µm.

**Movie S3**

Live cell imaging of Bax deficient HeLa, pTET3G-BNIP5(WT) and GFP-Bak(Green) expressing cells in medium. Mitochondria were labelled with Mitotracker deep red (Cyan). Time is shown as hh:mm:ss. Scale bar is 10 µm.

**Movie S4**

Live cell imaging of Bax deficient HeLa, pTET3G-BNIP5(WT) and GFP-Bak(Green) expressing cells treated with doxycycline at the start of the movie. Mitochondria were labelled with Mitotracker deep red (Cyan). Time is shown as hh:mm:ss. Scale bar is 10 µm.

**Movie S5**

Live cell imaging of Bax deficient HeLa, pTET3G-BNIP5(WT) and GFP-Bak(Green) expressing cells treated with cycloheximide at the start of the movie. Mitochondria were labelled with Mitotracker deep red (Cyan). Time is shown as hh:mm:ss. Scale bar is 10 µm.
